# The *Phaeodactylum tricornutum* Diaminopimelate Decarboxylase was Acquired via Horizontal Gene Transfer from Bacteria and Displays Substrate Promiscuity

**DOI:** 10.1101/2020.10.01.322594

**Authors:** Vincent A. Bielinski, John K. Brunson, Agnidipta Ghosh, Mark A. Moosburner, Erin A. Garza, Zoltan Fussy, Jing Bai, Shaun M.K. McKinnie, Bradley S. Moore, Andrew E. Allen, Steven C. Almo, Christopher L. Dupont

**Author notes:** These authors contributed equally to this work (Vincent A. Bielinski, John K. Brunson). The author(s) responsible for distribution of materials integral to the findings presented in this article in accordance with the policy described in the Instructions for Authors (www.plantcell.org) is: Christopher L. Dupont.

## Abstract

Diatoms are predicted to synthesize certain amino acids within the chloroplast, including L-lysine via a diaminopimelate-dependent pathway. Herein, we report that the model diatom, *Phaeodactylum tricornutum*, possesses a chimeric lysine biosynthetic pathway, which coalesces bacterial and plant genes, and is terminated by a chloroplast-localized diaminopimelate decarboxylase (DAPDC, *Pt*LYSA). We show that while RNAi ablation of *P*t*LYSA* is either synthetically lethal or concomitant with a slower growth rate, Cas9-mediated mutagenesis of *PtLYSA* results in recovery of heterozygous cells lines, suggesting that *PtLYSA* is an essential gene. Previously characterized DAPDCs are unique within the PLP-dependent decarboxylases where catalysis occurs at the D-stereocenter of the substrate and display a strict stereochemical preference for a (D,L)- or *meso*-substrate and not the D,D- or L,L-isomers of diaminopimelate (DAP) to synthesize L-lysine. Using decarboxylation assays and differential scanning calorimetry analyses, we validate that *Pt*LYSA is a *bona fide* DAPDC and uncover its unexpected stereopromiscuous behavior in substrate specificity. The crystal structure of *Pt*LYSA confirms the enzyme is an obligate homodimer in which both protomers reciprocally participate in the active site. The structure underscores features unique to the *Pt*LYSA clan of DAPDC and provides structural insight into the determinants responsible for the substrate-promiscuity observed in *Pt*LYSA.

## INTRODUCTION

Lysine plays a variety of important roles in biology across the tree of life. Besides its role as an essential amino acid required for protein synthesis, L-lysine can also be degraded into the lipid building block acetyl-CoA via lysine catabolism pathways in some organisms, including *E. coli*, *Arabidopsis* and humans (Knorr et al., 2018; Zhu et al., 2004; Chang, 1978). In Gram-positive *Staphylococcus* strains, L-lysine is found as a structural component of the peptidoglycan layer and the biosynthetic enzymes required for peptidoglycan production are involved in β-lactam resistance (De Lencastre et al., 1999). Two main routes for the *de novo* production of L-lysine exist in nature, although new and unique pathways are still being discovered in bacteria (Price et al., 2018). Higher fungi, euglenids and some bacteria produce L-lysine from homocitrate via the α-aminoadipate pathway (Xu et al., 2006; Kosuge et al., 1998). Land plants and most bacteria employ four biochemically distinct variations of the diaminopimelate (DAP) pathway to synthesize L-lysine from an aspartic acid precursor (Pratelli and Pilot, 2014; Kirma et al., 2012; Jander and Joshji, 2009; Hudson et al., 2005). In plants, the pathway includes an aminotransferase activity that produces L,L-DAP as an intermediate. L,L-DAP undergoes a subsequent epimerization to *meso*-DAP (D,L-DAP), the substrate for the final enzymatic step, which is catalyzed by diaminopimelate decarboxylase (DAPDC) to produce L-lysine (Bukhari and Taylor, 1971; White and Kelly, 1965; Dewey and Work, 1952).

DAPDC proteins are unique in the amino acid decarboxylase family as the only members to carry out decarboxylation exclusively at the D-stereocenter of the substrate. All DAPDC enzymes characterized thus far display a strict substrate preference for *meso*-DAP over other DAP stereoisomers and function as homodimeric complexes (Peverelli et al., 2016, Peverelli and Perugini, 2015, Griffin et al., 2012; Hu et al., 2008, Hutton et al., 2007). Elimination of CO_2_ from *meso*-DAP by DAPDC requires the cofactor pyridoxal-5’-phosphate (PLP), which forms a Schiff base with a conserved lysine residue (K54 in *E. coli* LYSA, UNIPROT ID P00861) in the active site of the enzyme (Ray et al., 2002). PLP facilitates the decarboxylation of *meso*-DAP substrate via formation of a Schiff base involving the ααamino group of the *meso*-DAP D-stereocenter, while distal residues in the DAPDC catalytic pocket interact with the L-stereocenter of *meso*-DAP (Hu et al., 2008, Gokulan et al., 2003). The structures of DAPDC orthologs reveal that both protomers of the dimer contribute to form the active sites in which a conserved arginine from the EPGR motif directs preference for *meso*-DAP (Crowther et al., 2019; Son and Kim, 2018; Weyand et al., 2009; Hu et al., 2008; Gokulan et al., 2003; Ray et al., 2002). These structures evince that DAPDC orthologs have a variable and flexible active site loop that shields the active site ligands from bulk solvent during catalysis (Hu et al., 2008).

In photosynthetic eukaryotes studied thus far, it appears the terminal steps of lysine biosynthesis occur exclusively within the chloroplast (Hudson, 2006; Perl, 1992; Wallsgrove, 1980). It has been postulated that lysine biosynthetic gene clusters from the cyanobacterial lineage were obtained via the primary plastid endosymbiotic event that resulted in the emergence of Archaeplastida that encompass glaucophytes, red and green algae and land plants (Reyes-Prieto, 2012; Kidron, 2007; Velasco, 2002). Diatoms are ubiquitous ocean phytoplankton and major contributors to the carbon cycle (Armbrust, 2009), which were derived from a secondary endosymbiosis event and thrive in upwelling-induced, nutrient-rich conditions making them the basis for the world’s shortest and most energy-efficient food webs. Similar to land plants and green algae, diatoms are capable of *de novo* synthesis of essential amino acids, including L-lysine via a predicted aminotransferase-dependent DAP pathway (Figure 1). Genomic reconstruction of pathways in diatoms suggests many amino acid biosynthetic genes are present in multiple copies and function within organelles, while a subset of biosynthetic genes is present as single copies (Bromke, 2013; Bowler et al., 2008).

**Figure 1.**
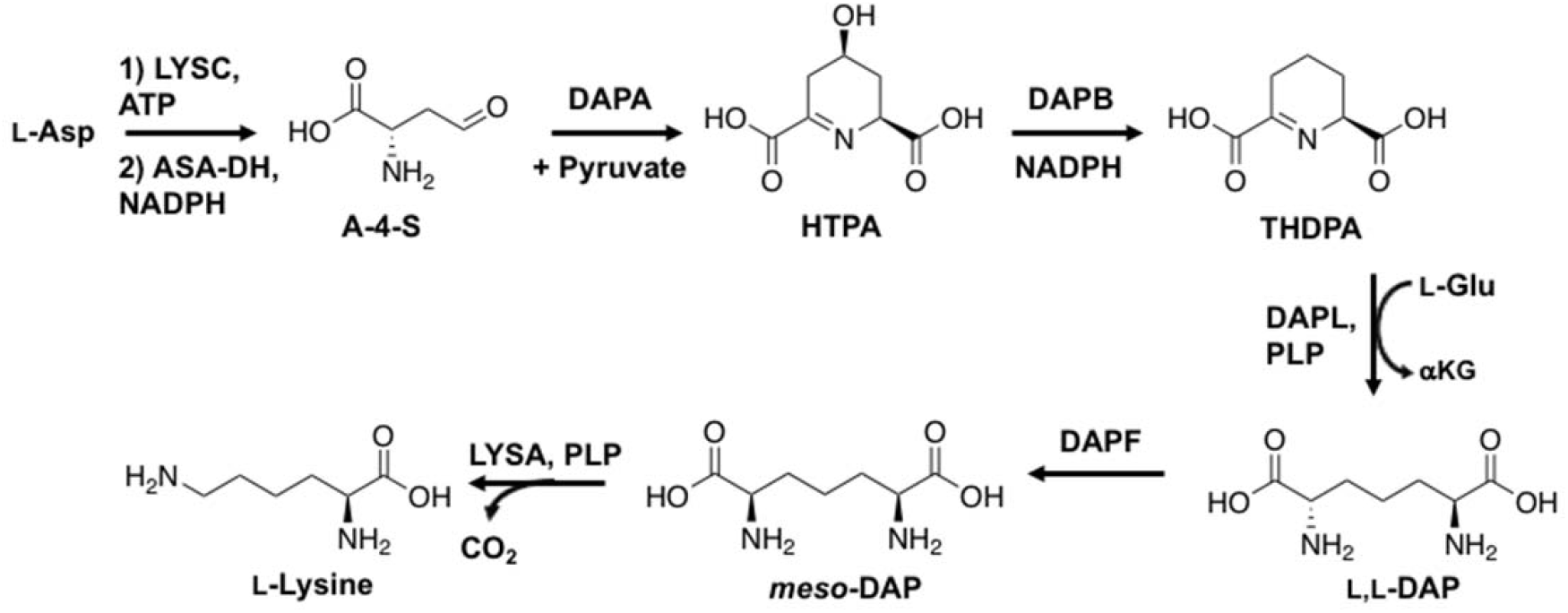
Lysine biosynthetic pathway in *P. tricornutum* based on predicted gene functions from the genome (Table 1). Entry of aspartate metabolites into the DAP pathway begins with addition of pyruvate to A-4-S via *DAPA* to produce 4-hydroxy-tetrahydrodipicolinate (HTPA), followed by *DAPB*-mediated reduction to THDPA (2,3,4,5-tetrahydrodipicolinate). DAP aminotransferase (*DAPL*) converts THDPA to L,L-diaminopimelate (L,L-DAP). L,L-DAP is subsequently isomerized to *meso*-DAP by *DAPF* epimerization activity, with lysine produced from *meso*-DAP via decarboxylation by *LYSA*

Within the genome of the model diatom *Phaeodactylum tricornutum*, the predicted lysine biosynthesis genes exist as single copies, and the corresponding proteins of the last four steps of the pathway predicted to localize within the chloroplast via canonical ASAF-type motifs and N-terminal chloroplast targeting peptides (Kilian and Kroth, 2005; Apt et al., 2002). In this report, we describe the phylogenetic origins of lysine biosynthetic machinery in diatoms. In addition, we report subcellular localization and biochemical activity of the enzyme involved in the terminal step of lysine biosynthetic pathway in *P. tricornutum*, and reveal that the enzyme (Phatr3_J21592, UNIPROT ID B7G3A2; *Pt*LYSA hereafter) is essential and harbors unexpected promiscuity in substrate utilization. In order to gain insight into this unique behavior, we determined the crystal structure of *Pt*LYSA, and defined the determinants responsible for substrate promiscuity in diatoms.

## RESULTS

### Phylogenetic Analysis

The DAP biosynthetic pathway in diatoms appears to be a hybrid of components from plants and bacteria potentially acquired via horizontal gene transfer (HGT), a process previously described in sequenced diatom genomes (Bowler, 2008). For the five enzymes catalyzing the synthesis of L-lysine from L-aspartate 4-semialdehyde (Table 1), orthologs in sequenced genomes and transcriptomes were identified, aligned, and examined using a maximum likelihood analysis (Figure 2). In parallel, target peptide analyses were conducted to predict subcellular localization (Table 1). The gene *PtDAPA* (Phatr3_J11151) encodes a putative dihydrodipicolinate synthase (DDPS) homolog in *P. tricornutum* and cannot be confidently localized based on target peptide analysis (Table 1). Our analyses reveal that *PtDAPA* has orthologs in other stramenopile genomes; however, the closest ancestors are in gammaproteobacteria (Figure 2A). For the three intermediate steps in the pathway, the diatom proteins (carried out by *PtDAPF, PtDAPL* and *PtDAPB*) also have orthologs in the stramenopiles, the closest ancestors of which are found in plant or cyanobacterial genomes (*DAPL* in Figure 2B, *DAPF and DAPB* in Supplemental Figure 1). Each of these proteins also contain putative N-terminal chloroplast targeting sequences (Gruber and Kroth, 2017; Apt et al., 2002).

**Table 1.** *P. tricornutum* lysine biosynthetic pathway protein IDs, phylogenetic outgroup, and predicted subcellular localization.

| **Pt Gene ID** | **Function** | **EC ID** | **Ancestor** | **Localization** |
| --- | --- | --- | --- | --- |
| Phatr3_J11151 | *DAPA*, 4-hydroxytetrahydropicolinate synthase | 4.3.3.7 | *Vibrio* | Unclear |
| Phatr3_J4025 | *DAPB*, 4-hydroxytetrahydropicolinate reductase | 1.17.1.8 | Plant | Chloroplast |
| Phatr3_J22909 | *DAPL*, L,L-diaminopimelate aminotransferase | 2.6.1.83 | Plant | Chloroplast |
| Phatr3_J34582 | *DAPF*, diaminopimelate epimerase | 5.1.1.7 | Plant | Chloroplast |
| Phatr3_J21592 | *LYSA*, diaminopimelate decarboxylase | 4.1.1.20 | *Verrucamicrobia* | Chloroplast |

The terminal step of the DAP pathway in *P. tricornutum* is predicated to be catalyzed by *PtLYSA* (Phatr3_J21592), which has orthologs throughout the stramenopile lineage; however, our analyses strongly suggest that the stramenopile lineage acquired the gene through horizontal gene transfer from Bacteria, specifically the PVC (Planctomycete-Verruca-Chlamydia) supergroup (Figure 2C). The diatom sequences for the enzymes in the pathway form monophyletic clades in phylogenetic inferences, suggesting that these enzymes are conserved through the entire lineage. In addition, our analyses reveal that the long branch length of the diatom and *Bolidophyceae LYSA* clade is associated with the insertion of a highly conserved 9 amino-acid segment into the active site loop of the predicted protein (Figure 2D). In this work we focused on experimentally validating the terminal step of this chimeric lysine biosynthetic pathway for the following reasons: **1**) bioinformatic annotations of the *P. tricornutum* genome suggest the presence of three putative organic acid decarboxylase genes (Bowler et al., 2008) however the LYSA enzymes in diatoms has not been biochemically validated thus far, **2**) the presence of the chloroplast targeting sequence in the *Pt*LYSA protein suggesting a similar localization of the pathway observed in plants, and **3**) inclusion of a highly conserved substitution in a critical substrate-binding residue and a unique insertion in the active site loop, while maintaining its ancestral link to bacteria in the PVC supergroup.

### Determination of *Pt*LYSA Subcellular Localization

The subcellular localization of *PtLYSA* in *P. tricornutum* was determined using transgenic and exconjugant diatoms expressing C-terminal fusions of *Pt*LYSA with either YFP or mTurquoise2 (mT2). Transgenic diatoms expressing *Pt*LYSA-YFP fusions driven by *pFCPB* promoter were generated by particle bombardment (Apt et al., 1996). Confocal microscopy showed that on occasion the YFP signal formed distinct punctate distributions within the chloroplast (Figure 3A), while in other transgenic strains the YFP signal was located throughout the chloroplast. Exconjugants expressing similar *Pt*LYSA-mT2 protein fusions on stable episomes were also generated, which provides a more controlled heterologous expression platform (Karas et al., 2015). Confocal microscopy of *Phaeodactylum* expressing a C-terminal mT2-fused version of *Pt*LYSA revealed a distinct regional overlap of the mT2 fluorescence with the autofluorescence of chlorophyll in the chloroplast (Figure 3C). We observed that exconjugant cells with constitutive expression of *PtLYSA*-mT2 showed signs of cell stress, grew poorly and adopted an ovoid morphology. We chose to employ the nitrate inducible pNR promoter (Chu et al., 2016; Poulsen and Korger, 2005) to repress *Pt*LYSA-mT2 expression during conjugation and strain selection and inducing expression after colony isolation. Exconjugants were generated and selected using ammonium as the nitrogen source and upon induction of cultures with nitrate, the cell morphology of these cultures quickly changed from pennate to ovoid and the cells began aggregating. In comparison, cultures expressing a cytosolic version of mT2 had little to no ovoid cells and grew to a higher cell density.

**Figure 3.**
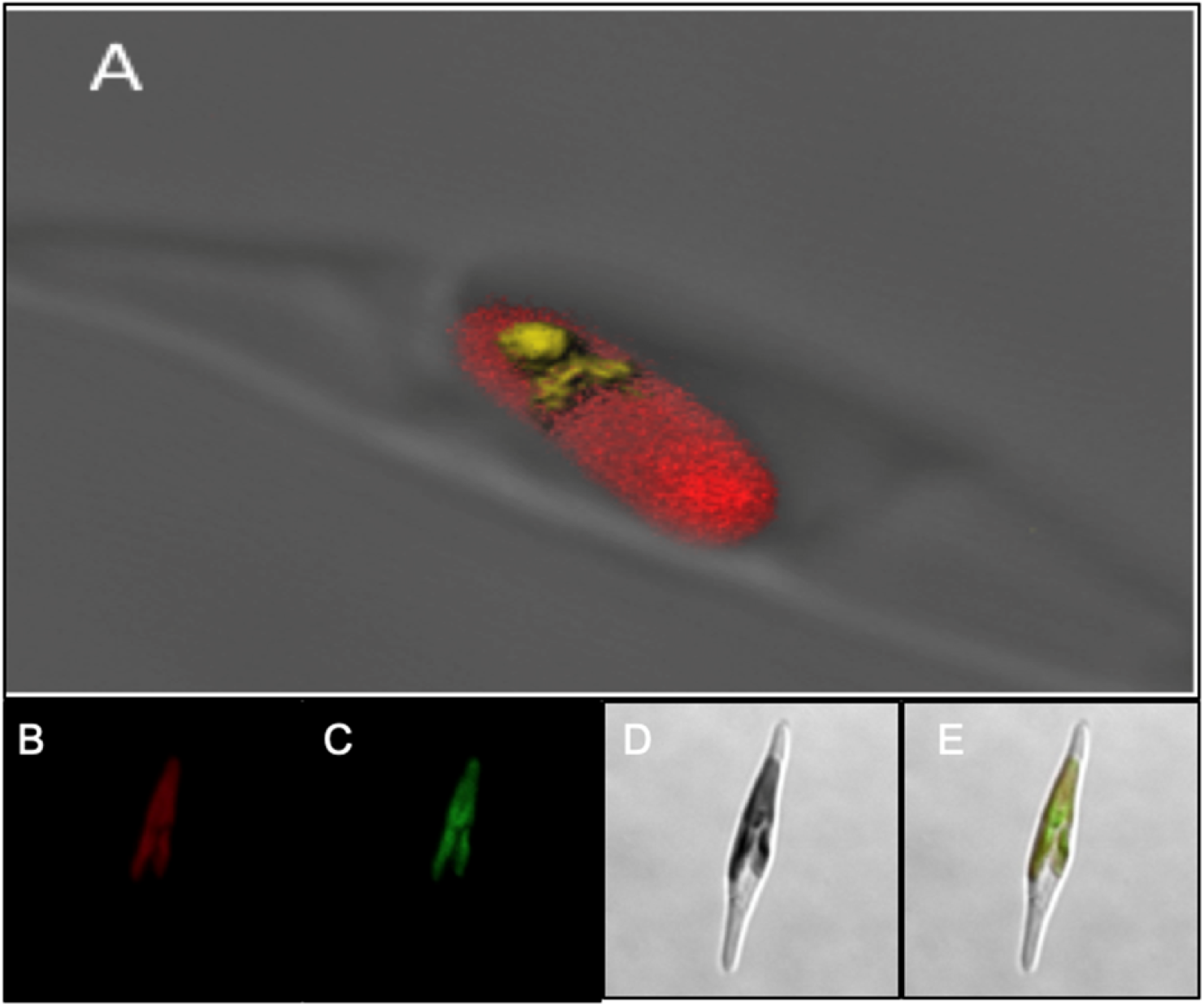
*In vivo* localization of C-terminal FP-tagged *Pt*LYSA constructs in *P. tricornutum.* **(A)** Z-stacked confocal microscopy image of *Pt*LYSA-YFP. Yellow, YFP signal; Red; Chlorophyll autofluorescence. **(B)** Confocal microscopy of *Pt*LYSA-mT2, chlorophyll autofluorescence channel. **(C)** Confocal microscopy of *Pt*LYSA-mT2, mTurquoise2 channel. **(D)** Confocal microscopy of *Pt*LYSA-mT2, bright field channel. **(E)** Merge of B,C,D

### *PtLYSA* RNAi and CRISPR

Reported transcriptomic analysis of the lysine pathway genes in *P. tricornutum* reveals the expression patterns of the associated genes are diel-regulated, with the aminotransferase *PtDAPL* showing the greatest change in expression over the course of the day (Smith et al., 2016; Supplemental Figure 2). In order to determine if the putative *Pt*LYSA protein is necessary for growth, we performed RNAi-driven gene knockdown, as well as TALEN and CRISPR-Cas9 gene mutagenesis experiments. For the RNAi-based gene knockdown, overlapping 250-bp and 400-bp intragenic fragments of *PtLYSA* were generated and cloned into a receiver vector as described (De Riso et al., 2008), with the exception that Gibson assembly (Gibson et al., 2008) was used instead of classical restriction enzyme dependent cloning. The receiver vector contains a transcriptional unit conferring phleomycin resistance and a cloning site with either the *pFCPB* or *pNR* upstream of the cognate terminator sequence. All attempts at attaining transformants of RNAi driven by *pFCPB* failed to yield colonies, despite the inclusion of a non-lethal positive control targeted to the *P. tricornutum* urease gene (Phatr3_J29702) that yielded transformants. We were able to attain RNAi-expressing transgenic lines where the hairpin loop for *PtLYSA* is driven by *pNR* only if the transformation is plated with ammonia as a sole nitrogen source. These lines exhibit reduced growth rates and *PtLYSA* protein content (Supplemental Figure 3) when grown on nitrate (RNAi-induced) instead of ammonia (RNAi-silenced).

Gene ablation vectors were made with the TALEN system described for *P. tricornutum* (Weyman et al., 2015); while colonies were obtained for the positive control urease knockout lines, none could be recovered for cell lines transformed with vectors targeting *PtLYSA*. Subsequent attempts were carried out using episome-delivered CRISPR-Cas9 methodology that includes multiplexed small-guide RNAs (sgRNAs, Moosburner et al., 2020). Two sgRNAs were delivered with Cas9 together or individually to *Phaeodactylum* using bacterial conjugation (Diner et al., 2016; Karas et al., 2015). The first attempt to knockout *PtLYSA* used a Cas9-fusion episome that harbors both sgRNAs, g21592-1 and g21592-2. Initially, two transformation attempts were made to produce transgenic *P. tricornutum* lines. Each time, less than 20 colonies were obtained on the plate, even after transformation optimization. On the third attempt, the selection plate growth media was supplemented with L-lysine (40 μg mL^−1^), which produced over 100 colonies on the selection plates.

A total of 24 colonies from the third conjugation attempt were analyzed for Cas9-induced indels by Sanger sequencing and TIDE analysis. The expected mutagenesis genotype was an 18-bp deletion between the two sgRNA cleavage sites. Manual curation of the sequences did not produce any obvious knock-out mutations but had a signature of Cas9 activity at the expected cut site(s). TIDE analysis revealed that 6 of 24 cell lines were heterozygous, or heterozygous where one allele contained an 18-bp deletion and the other allele retained a wild-type sequence (Figure 4). This observation was surprising since an 18-bp deletion should not knock-out the protein but could result in reduced enzyme activity or protein misfolding by removing 6 amino acids within the open-reading frame. Other attempts to produce a biallelic mutant cell line included re-streaking colonies to produce monoclonal cell lines and growing the heterozygous cell lines on increasing concentrations of L-lysine to force a biallelic mutation to occur. Nonetheless, the only genotype that could be produced using both sgRNAs to cut out an 18-bp sequence was heterozygous.

**Figure 4.**
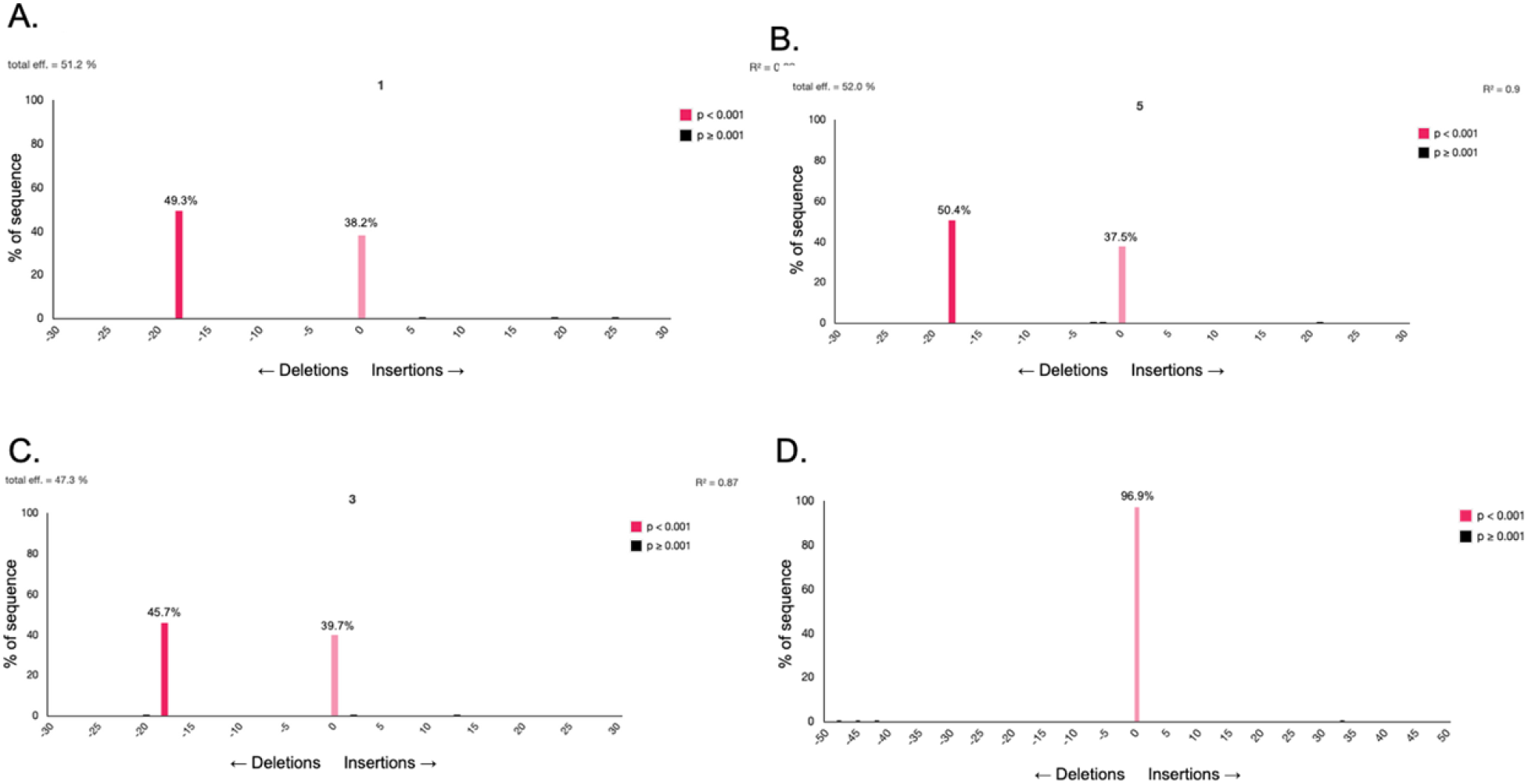
TIDE sequencing analysis results for heterozygous *P. tricornutum PtLYSA* mutants produced using two sgRNAs (gLysA-1 and gLysA-2). **(A-C)** Heterozygous mutation of an 18-bp deletion paired with a wild-type sequence for three cell lines. **(D)** TIDE result for a wild-type sequence of *PtLYSA*.

Subsequently, each sgRNA was delivered to *Phaeodactylum* individually and selected using growth media supplemented with L-lysine. 24 colonies were picked and analyzed for genotypes as described previously. Colonies that contained g21592-2 could not produce a transgenic genotype and were all wild-type at the *PtLYSA* locus. As for g21592-1, 2 of 24 colonies showed the effects of Cas9 activity after manual sequence curation and TIDE analysis. Since the induced mutations would be random indels, the colonies were re-streaked and monoclonal cell lines were picked again. Of 24 colonies, 18 had Cas9 activity at the g21592-1 cut site. TIDE analysis revealed that one third (8/24) of colonies were clearly heterozygous (Figure 4). Three colonies had the genotype of a 12-bp deletion plus wild-type and five colonies had the genotype of a 10-bp deletion plus wild-type. The other colonies had either a wild-type genotype (4), a mixed genotype without a wild-type signature (5), or a mix of indel and wild-type (2) (Supplemental Figure 4). In the 10-bp deletion plus wild-type heterozygous cell lines, a deletion of a 10-bp sequence would result in a deleterious frame-shift by introducing a premature stop codon in one of the two alleles. Therefore, one functional allele is sufficient to retain L-lysine biosynthesis in the chloroplast; however, using two sgRNAs, a biallelic mutant cell line of *PtLYSA* could not be obtained.

### Biochemical Analysis of *Pt*LYSA

Primary sequence analysis indicated that *PtLYSA* is homologous to an organic acid decarboxylase, likely a DAPDC, a well-studied and structurally characterized enzyme family (Hu et al., 2008; Gokulan et al., 2003; Ray et al., 2002). The *P. tricornutum* genome is predicted to harbor genes for putative DAP, ornithine and arginine decarboxylases (Smith et al., 2016; Bowler et al., 2008). To biochemically validate *Pt*LYSA as a DAPDC, we cloned full-length and varying N-terminal deletion constructs into a pBAD-driven bacterial expression vector, fusing them with a C-terminal 6xHis-Flag affinity tag (Savitsky et al., 2010). To screen for expression and solubility at small scale, the resultant plasmids were transformed into *E. coli* BL21 cells (Supplemental Figure 5). Overnight induction with 0.5% L-arabinose at 30 °C yielded several soluble constructs, with increased solubility at 18 °C. Judging by the level of expression and solubility, we chose to purify the Δ36-*Pt*LYSA-F481L variant protein in large quantities using immobilized metal (Ni) affinity chromatography (IMAC) and tested the purified enzyme for *in vitro* activity towards multiple substrates including ornithine and all three DAP stereoisomers (Figure 5, Supplemental Figure 6; Methods). The resolution of substrates and products of Δ36-*Pt*LYSA-F481L was achieved through pre-column derivatization with 1-fluoro-2,4-dinitrophenyl-5-L-alanine amide (L-FDAA), known as Marfey’s reagent (Bhushan and Brückner, 2004).

**Figure 5.**
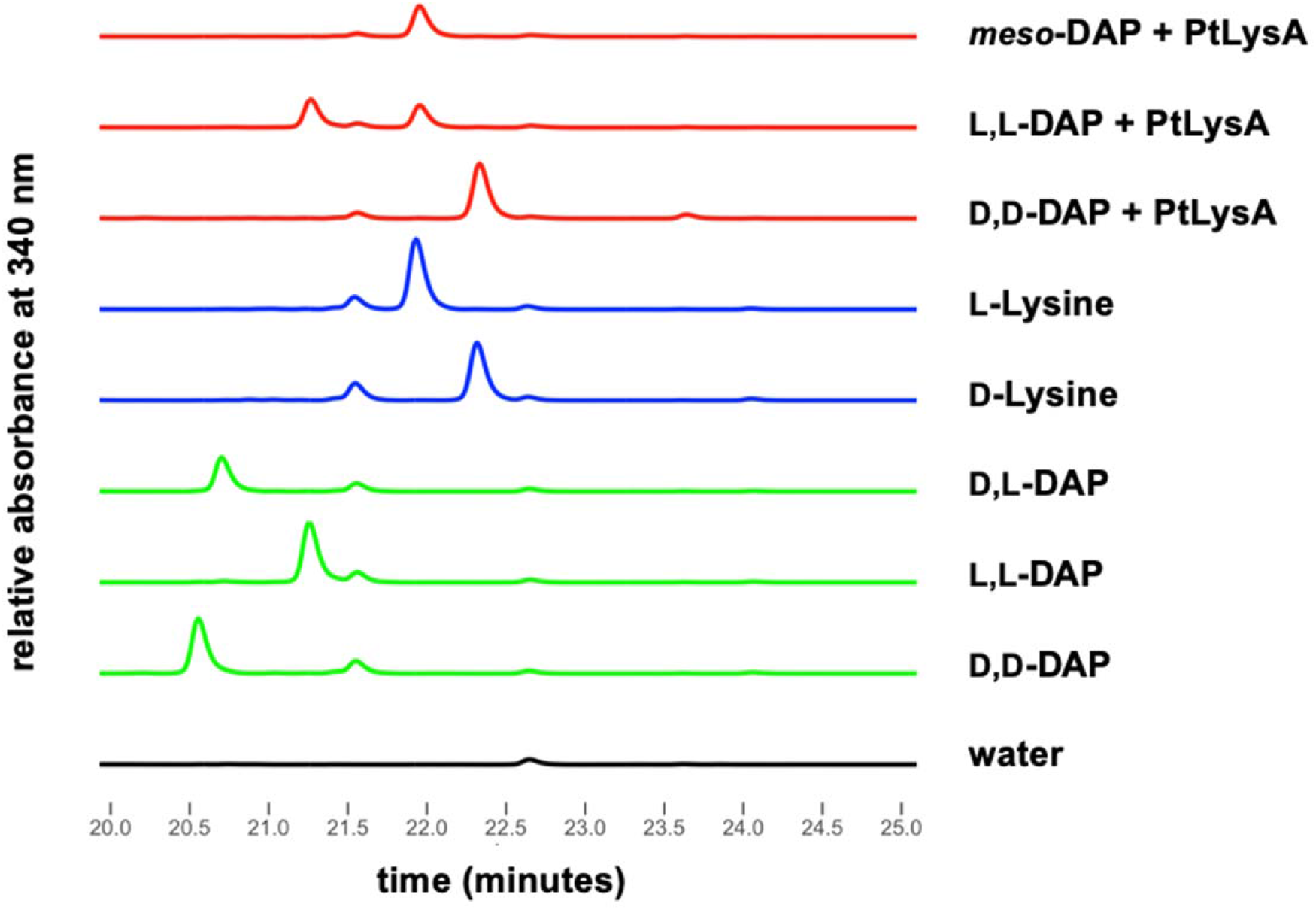
RP-HPLC (λ = 340 nm) analyses of L-FDAA (Marfey’s) derivatized *Pt*LYSA reactions with *meso*-, L,L-, and D,D-DAP, and comparison to similarly derivatized lysine and DAP standards. Substrates for enzyme reactions were added at 1 mM concentration and similarly 1 mM of each standard was used for subsequent derivatization. Overnight *Pt*LYSA reactions of *meso*-DAP and D,D-DAP produced L- and D-lysine, respectively. L,L-DAP reactions also produced L-lysine, but did not proceed to completion.

**Figure 6.**
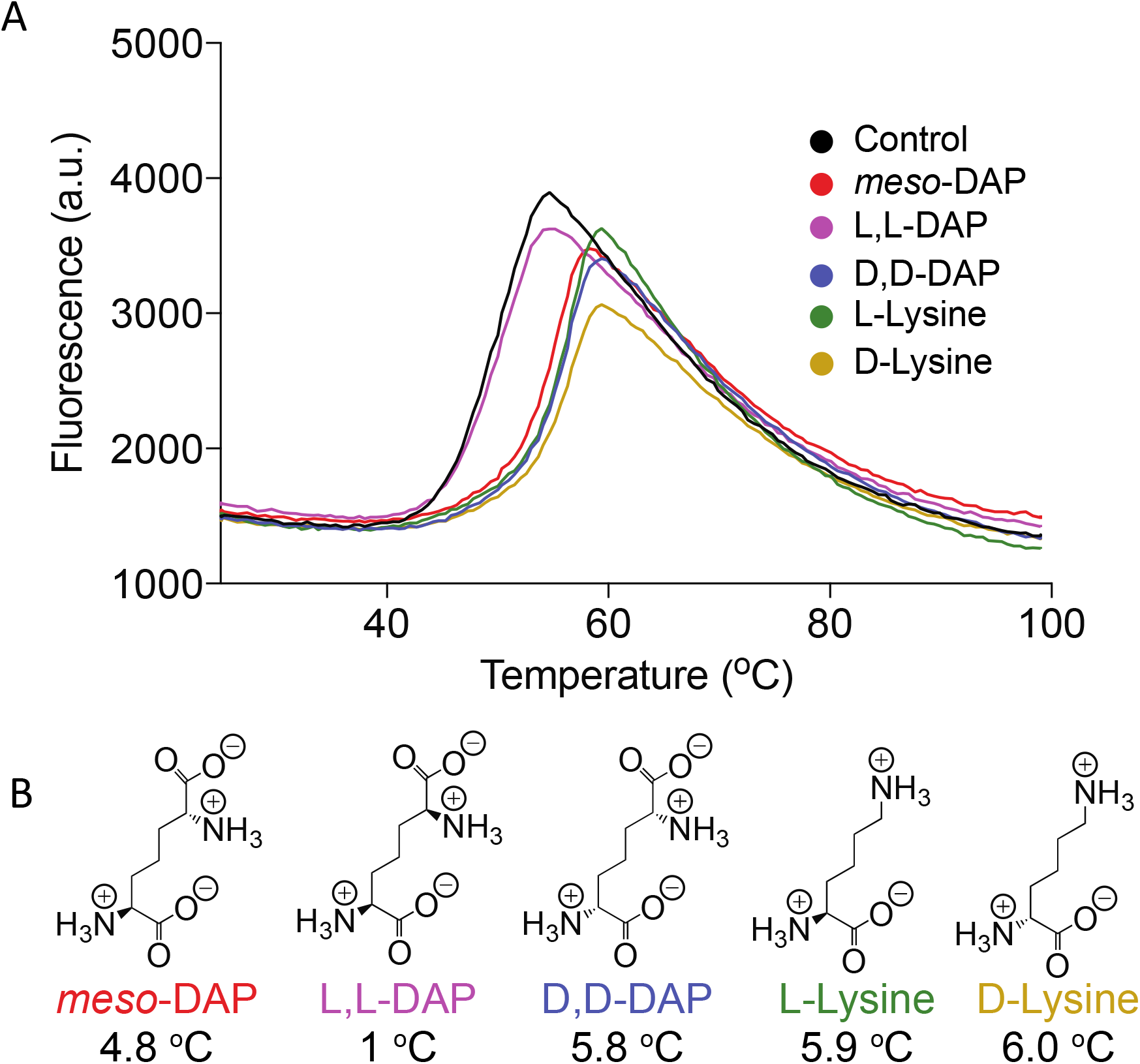
Differential scanning fluorimetry of *Pt*LYSA in presence of potential ligands. **(A)** Denaturation of *Pt*LYSA as a function of temperature as observed by increase in fluorescence of the indicator dye SYPRO Orange, which binds nonspecifically to hydrophobic surfaces. At higher temperatures, the intrinsic fluorescence degrades due to the formation of protein aggregates and dye dissociation. **(B)** Schematic description of ligands and their calculated ΔT_m_ values are shown.

Derivatization of DAP and lysine with Marfey’s reagent allows for chromatographic separation of stereoisomers followed by UV-detection (340 nm) and was used for reaction assay analysis by HPLC (Methods). We found that the purified *Pt*LYSA enzyme produced L-lysine from *meso*-DAP and displayed no activity towards ornithine (Supplemental Figure 6), confirming biochemical function of *Pt*LYSAas a DAPDC. When the purified Δ36-*Pt*LYSA-F481L protein was challenged with three different DAP stereoisomers, to our surprise, we observed that the enzyme is capable of producing D-lysine from the D,D-DAP and L-lysine from L,L-DAP. Both the *meso-*DAP and D,D-DAP reactions go to completion; however, the L,L-DAP reaction has significant amount of unreacted substrate even after overnight incubation (Figure 5).

*Pt*LYSA was also subjected to differential scanning fluorimetry (DSF) against a pool of potential ligands (Figure 6). Purified *Pt*LYSA exhibited a melting temperature (T_m_) of 49 °C (± 1) and is stabilized by the potential ligands. The substrate *meso-*DAP increased thermal stability (ΔT_m_) of *Pt*LYSA by 4.8 °C (± 0.8), indicating a potential interaction between *meso*-DAP and *Pt*LYSA. Similarly, *Pt*LYSA was stabilized by the product, L-lysine (ΔT_m_= 5.9 °C (± 1)). DSF data revealed that the D,D-DAP isomer was able to stabilize *Pt*LYSA by 5.8 °C (± 0.6); however, L,L-DAP enhanced the thermal stability by 1 °C (T_m_= 50 °C (± 0.2); Figure 6). These DSF experiments are consistent with the preferred utilization of *meso*-DAP and D,D-DAP as substrates, relative to L,L-DAP. These data are also consistent with the incomplete processing of L,L-DAP observed during *in vitro* assays (~50% turnover to L-lysine after overnight incubation, Figure 5), in contrast to using D,D-DAP and *meso*-DAP to yield D-lysine and L-lysine, respectively. In addition, D-lysine binds to *Pt*LYSA and induces a similar thermal stability similar to L-amino acid (Figure 6), suggesting that the catalytic pocket of *Pt*LYSA accommodates both enantiomeric forms of lysine.

The use of D,D-DAP as a substrate and the production of D-lysine is remarkable, as reported DAPDC orthologs, thus far, exhibit strong selectivity towards *meso*-DAP as substrate. Structural analyses reveals that amino acids sidechains in DAPDC active sites orient *meso*-DAP for catalysis and restrict D,D-DAP from engaging in a productive orientation with respect to the bound PLP cofactor necessary for decarboxylation reaction (Hu et al., 2008; Gokulan et al., 2003; Ray et al., 2002). In particular, a conserved arginine side chain in DAPDC orthologs appears to impose extensive steric hindrance to bar D,D-DAP; interestingly in diatoms the arginine side chain at this position is replaced by a threonine (T317 in *Pt*LYSA).

### Crystal Structure of *Pt*LYSA

On the basis of reported coordinates of DAPDC enzymes in the RCSB Protein Data Bank, a construct of *Pt*LYSA encompassing amino acids 39-476, which lacks the N-terminal signal peptide, was purified and co-crystallized in the presence of D-lysine (Methods). The structure of *Pt*LYSA was determined by molecular replacement using *E. coli* DAPDC (PDB: 1KNW) as a search model. The final model refined at 2.78 Å resolution had R_cryst_ and R_free_ values of 0.18 and 0.23, respectively (Table 2).

**Table 2.**
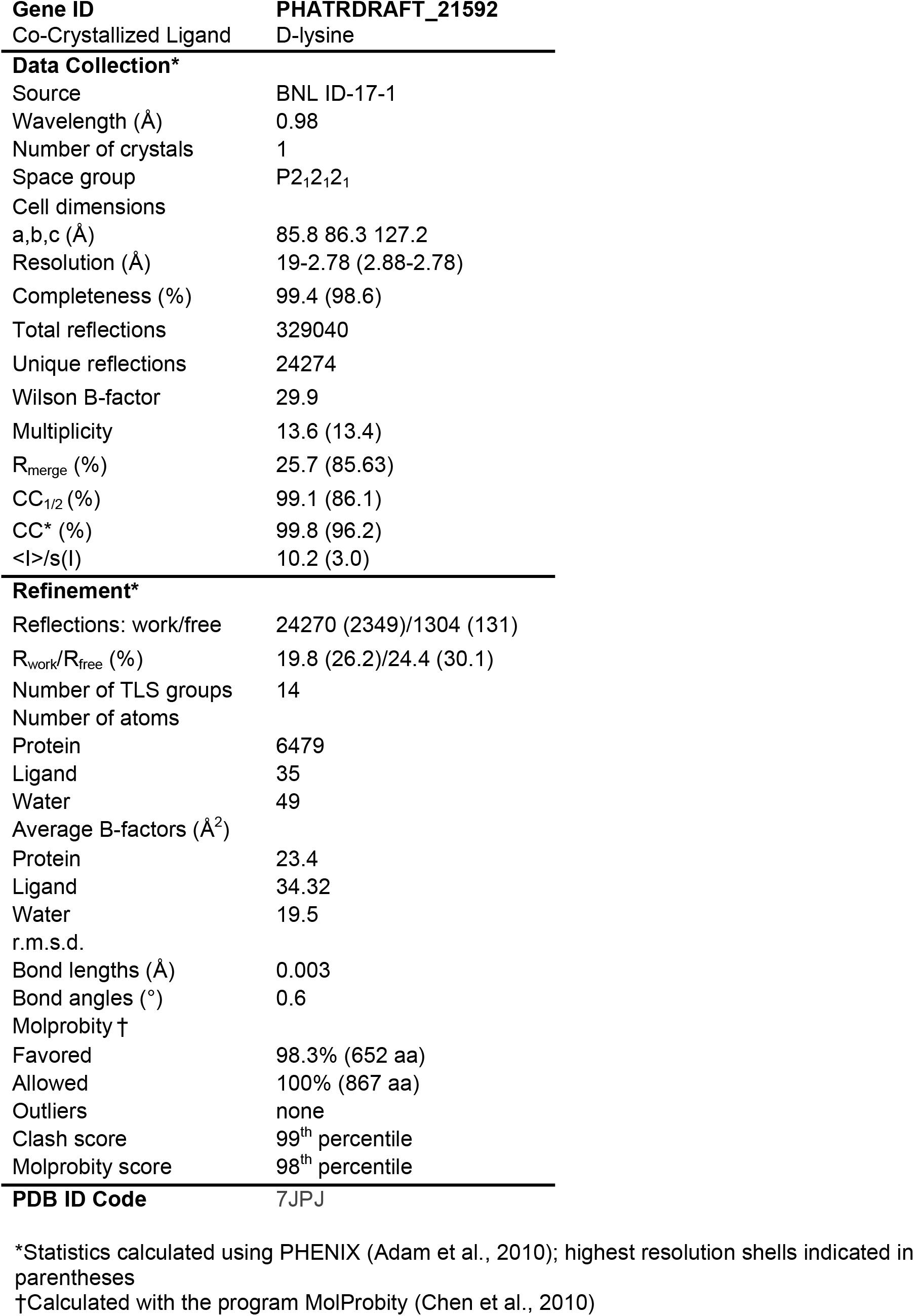
Crystallographic data collection and refinement statistics

Crystals contained two molecules of *Pt*LYSA in the asymmetric unit (protomers 1 and 2; Figure 7A), suggesting a dimer. Size-exclusion chromatography is consistent with the proposed dimer (elution time corresponding to an apparent molecular weight of ~100 kDa; monomer molecular weight of ~50 kDa) (Supplemental Figure 7), suggesting the protein is a dimer in solution. The two protomers are arranged in a “head-to-tail” configuration related by 2-fold non-crystallographic symmetry. The interface is polar in nature and shows remarkable surface complementarity while burying a total of ~4120 Å^2^ of solvent assessable surface area (Supplemental Figure 8) upon dimerization.

**Figure 7.**
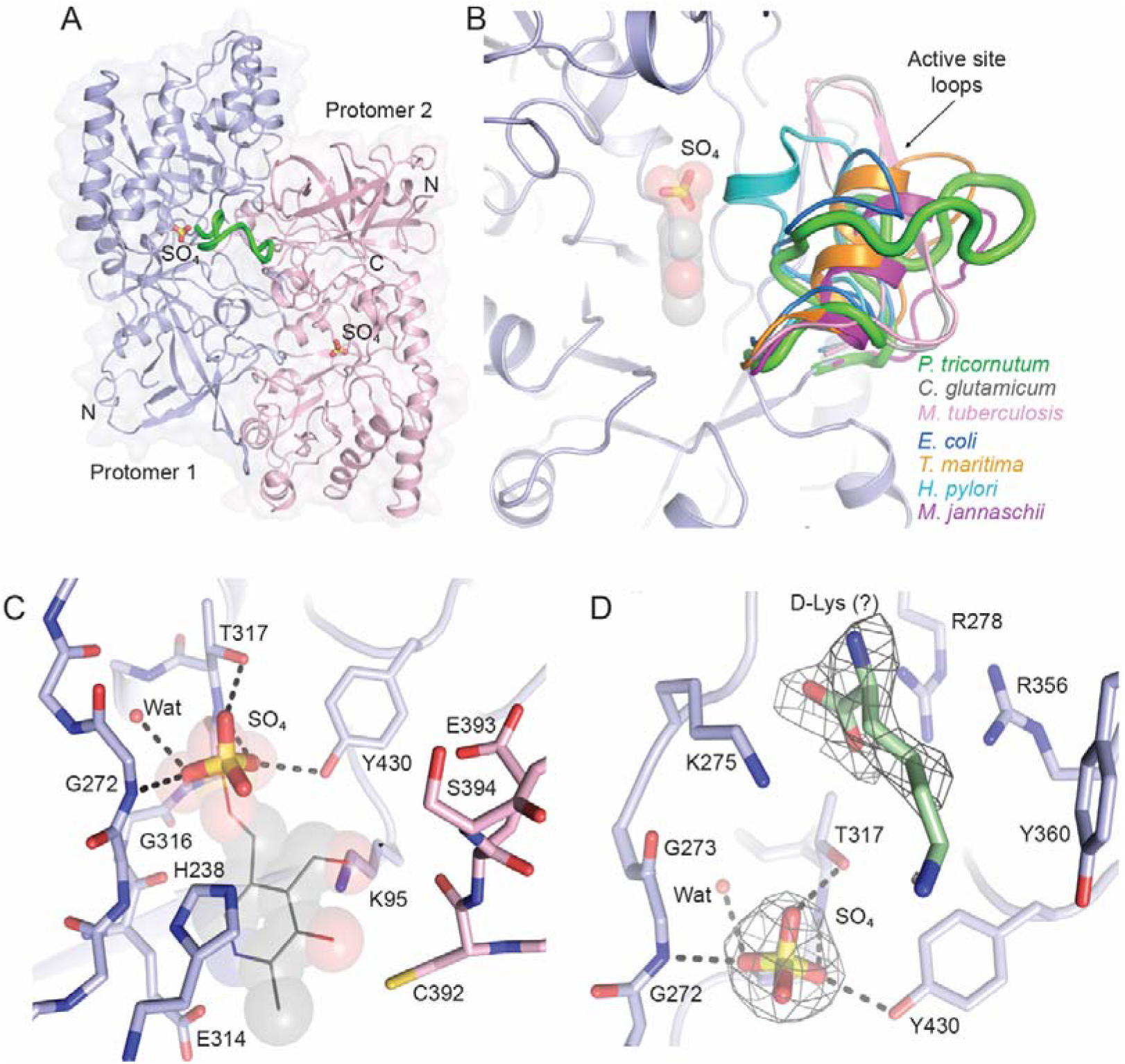
Structure of *Pt*LYSA highlighting the active site. **(A)** View of the *Pt*LYSA dimer shown in ribbon representation with secondary structure elements indicated by arrows (β strands) and ribbons (helices). The two protomers are colored in light-blue (protomer 1; chain A) and light-pink (protomer 2; chain B). A transparent molecular surface envelops the structure. The sulfate ions that occupies the active sites of the protomers are shown in stick representation and labeled. The ordered active site loop in protomer 1 is colored in green. N and C denote the location of the N and C termini. **(B)** Closeup view of the part of the active site in protomer 1 highlighting the bound SO_4_ and the active site loop (color coded as in A). Relative locations and structural elements of active site loops in *C. glutamicum* (PDB: 5X7M, in grey), *M. tuberculosis* (PDB: 2O0T, in pink), *E. coli* (PDB: 1KNW, in dark-blue), *T. maritima* (PDB: 2YXX, in gold), *H. pylori* (PDB: 2QGH, in cyan) and *M. jannaschii* (PDB: 1TUF, in magenta) DAPDC orthologs by aligning the respective coordinates on protomer 1. PLP (transparent spheres) was modeled into the protomer 1 based on its position in the *T. maritima* structure to denote the potential location and orientation of the co-factor in the *Pt*LYSA active site. **(C)** Closeup view and interactions of the SO_4_ in the active site in *Pt*LYSA. Contributing sidechains to the active site form the protomers (color coded as A) are shown in sticks and labeled. Contacts to SO_4_ are shown as dashed lines. Modeled PLP is shown in grey lines and transparent spheres. **(D)** Closeup view of the active site in *Pt*LYSA showing electron density from omit maps (contoured at 1.2σ; light-blue mesh) around the SO_4_ and putative D-lysine (stick representation, in green). Atomic contacts in *Pt*LYSA sidechains with SO_4_ are indicated by dashed lines. Potential sidechain contacts to D-lysine in the active site are shown stick representation. All figures depicting structure were generated with PyMol (DeLano et al., 2002).

**Figure 8.**
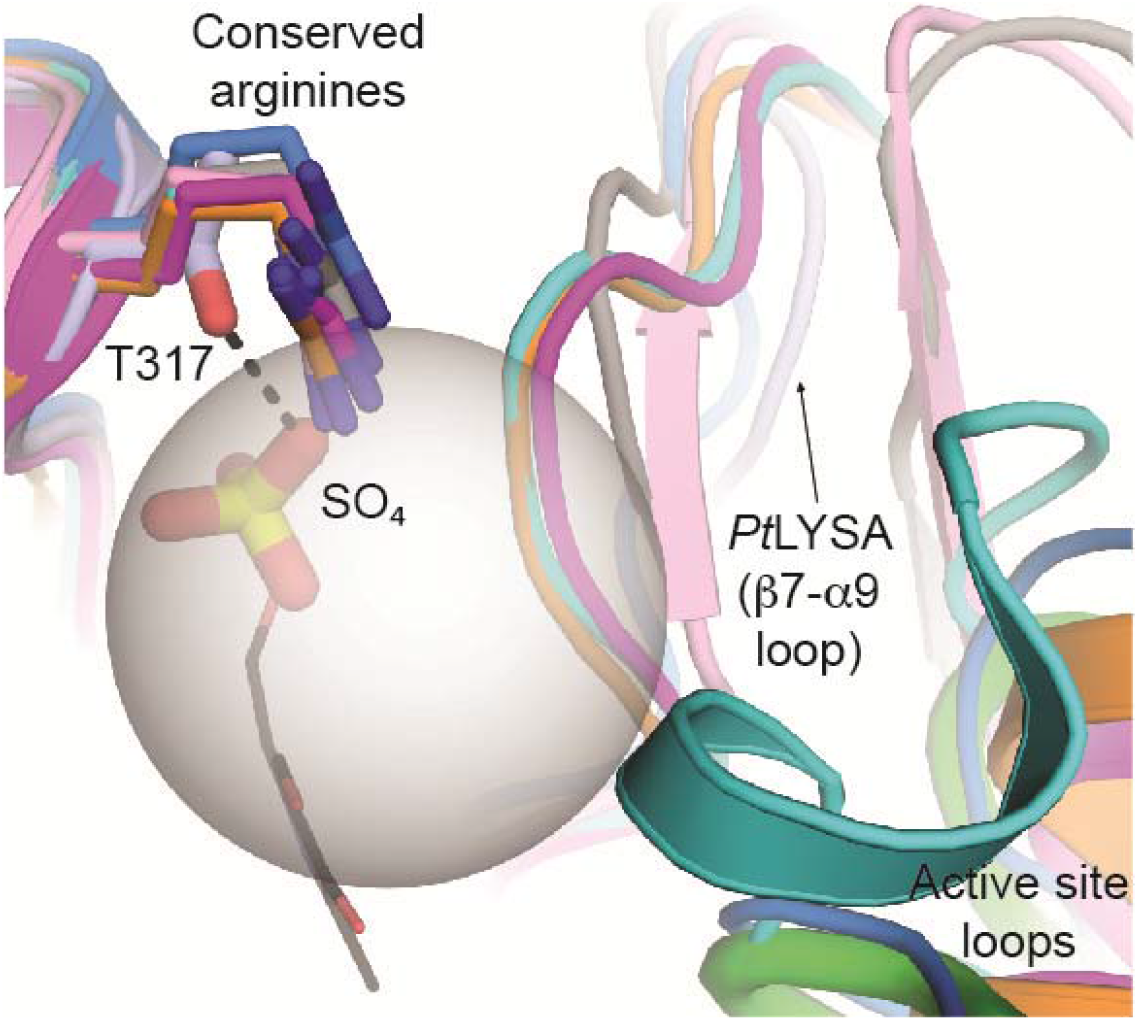
Distinctive active site in PtLYSA. The conserved arginine in DAPDC orthologs is replaced by a threonine, Thr317, (indicated in stick representation and color coded as in Figure 6) in *Pt*LYSA. SO_4_ in the active site is shown in stick representation. The connecting loop between the stand β7 and helix α9 (amino acid 236-244), which forms a wall of the active site, is smaller in comparison to the orthologs with exception of *E. coli* DAPDC (see Figure S8). The smaller sidechain of Thr317 together with the smaller β7-α9 connective loop adopting a distinct conformation (indicated by an arrow) compared to its counterpart create a distinctive and larger *Pt*LYSA active site (indicated by a transparent sphere) that can potentially accommodate unique substrate (D,D-DAP) for catalysis and can be exploited as specific druggable packet. Part of the active site loops in the orthologous DAPDC are shown and color coded as in Figure 6 and labeled. Modeled PLP from *T. maritima* (PDB: 2XYY) is shown in grey lines.

### Structure of *Pt*LYSA in relation to DAPDC orthologs

Individual protomers of *Pt*LYSA consist of two domains: an N-terminal 8-fold α/β barrel and a C-terminal β sandwich domain (Figure 7A), with both domains contributing to the dimer interface. The barrel domain lacks the N-terminal β sheet extension of *C. glutamicum* (Son and Kim, 2018), *M. tuberculosis* (Weyand et al., 2009) and *M. jannaschii* (Ray et al., 2002) enzymes; however, the domain is topologically similar to *H. pylori* (Hu et al., 2008), *E. coli* (PDB: 1KNW) and *T. maritima* (PDB: 2YXX) DAPDC, which lack the N-terminal extension. The protomers of *Pt*LYSA are similar to these last three DAPDC orthologs with a root-mean-square deviation (RMSD) between 1.1 and 1.3 Å over 249 aligned Cα atoms. Therefore, the structure unambiguously confirmed that *Pt*LYSA belongs to the group IV pyridoxal-5’-phosphate (PLP) dependent decarboxylase family (Kern et al., 1999; Sugio et al., 1995).

Structure-based sequence alignment of the DAPDC orthologs underscore two structurally divergent regions in *Pt*LYSA: **1**) the loop segment between β6 and α7 (β6-α7 loop; amino acids 183-209), and **2**) the loop segment between β7 and α9 (β7-α9 loop henceforth; amino acids 238-245) (Supplemental Figure 9). In *Pt*LYSA, the β6-α7 loop has a 7 amino acid insertion, which could only be built completely as a random coil into the electron density of protomer 1 (Figures 7A and 7B). Akin to protomer 2 of *Pt*LYSA, this segment is not included in the structures of DAPDC from *Aquifex aeolicus* (PDB 2P3E) and *Brucella melitensis* (PDB 3VAB). In the structure of *H. pyroli* DAPDC (PDB: 2QGH), the analogous segment has been annotated as ‘active site loop’, which occludes the active site ligands from solvent (Hu et al., 2008). In *H. pylori* this region forms a 3_10_ helix (Figure 7B). In *C. glutamicum* (PDB: 5X7M, Son and Kim, 2018) and *M. tuberculosis* (PDB: 2O0T, Weyand et al., 2009; PDB: 1HKW, Gokulan et al., 2003), this segment forms a two stranded anti-parallel β sheet (Figure 7B). In *M. jannaschii* (PDB: 1TUF, Ray et al., 2002) and *T. maritima* (PDB: 2YXX), this segment adopts a helical conformation; and in *E. coli* (PDB: 1KNW) it forms a similar, but smaller random coil compared to *Pt*LYSA segment (Figure 7B).

The β7-α9 loop forms a wall of the active site in all DAPDCs (Figure 8), and the *Pt*LYSA enzymes has a 5 amino acid deletion in this segment (Supplemental Figure 9), which causes an expansion of the *Pt*LYSA active site (Figure 8) compared to orthologs from *C. glutamicum*, *M. tuberculosis*, *M. jannaschii*, *H. pylori* and *T. maritima*. An analogous deletion has also been observed in the β7-α9 loop of *E. coli* DAPDC (Supplemental Figure 9), in which it adopts a conformation similar to that observed in *Pt*LYSA (Figure 8).

### Architecture of the substrate binding site in *Pt*LYSA

The structural analyses reveal that both protomers of the *Pt*LYSA dimer contribute to the active site by reciprocally sharing 8 conserved amino acids: Lys95, His238, Glu314, Tyr430, Cys392, Glu393, Ser394, and Thr317. The Thr317 is unique to *Pt*LYSA (Figure 7C, Supplemental Figure 9), as the corresponding residue in all structurally characterized DAPDCs is an arginine, which is part of the EPGR motif (Supplemental Figure 9) (Son and Kim, 2018, Weyand et al., 2009; Hu et al., 2008; Gokulan et al., 2003; Ray et al., 2002). While the sidechains of Lys95, His238, Tyr430 and Thr317 are supplied in *cis* to the active site of the protomers, the side chain of Cys392, Glu393 and Ser394, which form a conserved CESG/SD motif in DAPDC, are shared in *trans* from the other protomer (Figures 7C and 7D). In DAPDC orthologs, the conserved side chain at Lys73 from the Y/FAxKA motif (Supplemental Figure 9) forms a Schiff base with the cofactor, PLP, which exhibits aromatic stacking interaction with the side chain of His238. The side chains of Glu314 (OE) and Tyr430 (OH) contact (H-bonds) the pyridine endocyclic nitrogen (N1) and phosphate (O1P) of the PLP, respectively.

In the active sites of the *Pt*LYSA dimer, a sulfate ion occupies the position of the phosphate group of PLP (Figures 7C and 7D). Superimpositions of *Pt*LYSA protomers with PLP-bound DAPDC structures reveal that the coordination sphere of the sulfate ion is very similar to that observed in the PLP-PO_4_ interaction. However, there is a striking adaptation observed in the polyanion coordination in *Pt*LYSA; the sidechain oxygen (OG1) of Thr317 (EPGR motif; Supplemental Figure 9) is engaged in a H-bond interaction with the SO_4_ (Figures 7C and 7D). The arginine sidechains in DAPDC orthologs do not participate in such an interaction (Figure 8), instead they contribute to substrate specificity and stabilize the active site by interacting with a tyrosine sidechain OH via a H-bond. The tyrosine is conserved in DAPDC orthologs; however, it is replaced by an arginine, Arg278, in *Pt*LYSA.

Additional, but weak, electron density features were observed adjacent to the SO_4_ ion in the active sites of *Pt*LYSA. The densities were fenced by Lys275, Arg278, Arg358 and Tyr360, and interpreted and modeled as co-crystallized D-lysine for protomer 1 (Figure 7D). Given the weak density and modest 2.78 Å resolution of the refined coordinates, the assignment of D-lysine remains speculative. It is, however, notable that only Arg358 and Tyr360 are conserved among the DAPDC orthologs (Supplemental Figure 9).

## DISCUSSION

Diatoms evolved via serial endosymbiotic events between a photosynthetic symbiont and a non-photosynthetic exobiont that have occurred over the past 1.8 billion years (Armbrust et al., 2004). In addition to these processes, there have been substantial horizontal gene transfer events between diatoms and bacteria (Mock et al., 2017; Raymond et al., 2012; Bowler et al., 2008). The phylogenetic analyses reported within conclusively show the lysine biosynthetic pathway in diatoms is conserved across all species examined by genome or transcriptome sequencing to date. This conserved pathway includes genes likely derived from the symbiont in the primary endosymbiotic event (*DAPL, DAPB, DAPF)* as well as two genes (*DAPA, LYSA*) likely derived from cryptic horizontal gene transfers from Bacteria. *DAPA* appears to have originated from gammaproteobacteria in the *Vibrio* lineage, which is not a common Bacterial HGT partner for diatoms (Bowler, et al. 2009; Bowler et al., 2008). In contrast, the *LYSA* appears to have been transferred from the Verrucamicrobia lineage, which has been predicted to have contributed a substantial number of genes to modern diatom genomes (Bowler et al., 2008). The phylogenetically chimeric pathway is reminiscent of the urea cycle and overall nitrogen assimilation pathways within diatoms (Smith et al., 2019; Allen et al., 2011), both of which are composed of genes from different evolutionary partners of the serial endosymbiotic events. The conservation of this chimeric pathway across all diatoms sequenced to date suggests strong selective pressure for the retention of the pathway and that this event occurred early in the evolution of the diatom lineage.

Subsequent work focused on the terminal step of the pathway, which was hypothesized to be catalyzed by *Pt*LYSA. Biochemical assays confirmed the purified protein converts *meso*-DAP to L-lysine. The observation that overexpression of FP-linked fusions of *Pt*LYSA in *P. tricornutum* cells results in the co-localization of signal with chloroplast autofluorescence is consistent with the prediction that *PtLYSA* contains a canonical bipartite transit peptide harboring an ASAF cleavage site (Gruber and Kroth, 2017; Apt et al., 2002). This localization also corroborates previous reports that in plants lysine production is carried out in the chloroplast (Mills et al., 1980; Wallsgrove and Mazelis, 1980) and that the three genes upstream of *PtLYSA* (DAPF, DAPL and DAPB) encoding the terminal arm of the *P. tricornutum* lysine pathway also contain predicted N-terminal chloroplast targeting peptides (Table 1). The localization of these enzymes in the chloroplast has potential biochemical and system-wide consequences. These steps of the pathways will be subject to large changes in redox state (Rosenwasser et al., 2014; Baier and Dietz, 2005). Diurnal swings in chloroplast redox state may have significant implications for *DAPF*, where a dimerization is not only necessary for function but also sensitive to reversible disulfide-bond formation between protomers (Sagong and Kim, 2017). It is possible that redox-responsive disulfide bond disruption deactivates *DAPF* during the dark periods where the local chloroplast environment becomes more reducing, potentially halting the supply of *meso*-DAP. This potential post-translational effect is an additional level of regulation, superimposed upon that of diel gene expression of the entire lysine pathway, which is transcribed at higher levels during the light period in *P. tricornutum* (Smith et al., 2016) and may contribute to regulation of carbon flux and assimilation during the day/night cycle.

The dynamic regulation of the lysine biosynthetic pathway genes at the transcript level suggests an important role for this pathway in diatom metabolism. This notion is supported by the observed changes in the engineered gene expression during the course of this study. Overexpression of a heterologous copy of *PtLYSA* from an episome using a non-diel responsive promoter in a wild-type background induces a change in *P. tricornutum* cells from fusiform to ovoid morphotype, which is associated with cell stress (Lauritano et al., 2015). Controls show that this effect is not due to the expression of fluorescent tag. Similarly, homozygous gene knockouts were not attainable, with only heterozygous exconjugants being attained in media supplemented with exogenous L-lysine when Cas9 editing of *PtLYSA* was attempted. These data demonstrate a functional allele of *Pt*LYSA is essential for growth. These heterozygous exconjugants contained alleles coding for a native copy of *PtLYSA* as well as a copy with a 6 amino-acid deletion that is predicted to have minimal effect on the tertiary structure of the homodimeric enzyme. Deletion analyses identified the LEEAA sequence segment (amino acids 72-76) in *Pt*LYSA as essential (Supplemental Figure 9). This segment is not conserved among DAPDC orthologs, hence the basis for its essentiality was not immediately apparent. In the crystal structure, this segment forms a helix (α2 in *Pt*LYSA, Supplemental Figure 9), which is structurally conserved across the orthologs despite the lack of sequence similarity. The *Pt*LYSA structure underscores that this segment plays a secondary but important role in catalysis by buttressing residues in and adjacent to the active site (Supplemental Figure 10). We surmise that analogous deletion in DAPDCs would result in similar phenotypes in the orthologs. RNAi with constitutive promoters was also unsuccessful, though inducible RNAi successfully generated transgenic lines, which were used to verify that knockdown of *PtLYSA* expression has a detrimental effect on growth (Supplemental Figure 4). Overall, *PtLYSA* appears to be an essential gene in *P. tricornutum* and these results suggest that a diverse gene editing tool kit beyond just CRISPR-Cas guided gene knockouts is necessary to characterize *LYSA* and possibly other essential genes in diatoms.

The activity of *Pt*LYSA towards both *meso*-DAP and D,D-DAP (Figure 5) was unexpected, and is consistent with DSF data showing that *Pt*LYSA is able to bind both *meso*-DAP and/or D,D-DAP with similar ΔT_m_s (Figure 6). It should be noted that while the biochemical assay demonstrates that *Pt*LYSA can use L,L-DAP as substrate, and the reaction does not proceed to completion under the conditions employed (Figure 5). Additionally, L,L-DAP did not impact the thermal stability of *Pt*LYSA (Figure 6). Metabolomic studies in diatoms are in their infancy, therefore it is not known if diatoms produce D,D-DAP *in vivo*, or encounter it in its environment. Production of D,D-DAP *in vivo* would also be biochemically unprecedented, as D,D-DAP has never been demonstrated to be a physiological substrate for DAPDC enzymes, nor has it been demonstrated to be a potential product of DAPF epimerase activity in any organism (Hor et al., 2013). Therefore, the topic remains an open area of investigation. However, flexibility in substrate utilization may be advantageous under selective pressure, allowing organisms to take advantage of non-canonical substrates or cofactors encountered in the environment.

The structure of *Pt*LYSA confirms that the protein forms a “head-to-tail” symmetric homodimer, and buries a large amount of solvent accessible surface area, including the active site (Supplemental Figure 8). The extensive dimer interface of *Pt*LYSA along with the *cis-trans* orientation of the active site and its behavior in solution (Supplemental Figure 7) suggests that the protein is an obligate dimer, consistent with orthologs from diverse organisms (Son and Kim, 2018; Peverelli et al., 2016; Griffin, 2012 et al.; Weyand et al., 2009; Hu et al., 2008; Gokulan et al., 2003; Ray et al., 2002). In this aspect, *Pt*LYSA is similar to members of the DAPDC superfamily; however, unique aspects of the structure also provide some insight as to the stereochemical origins of the stereopromiscuity. The active site of *Pt*LYSA possesses three striking adaptations: **1**) an insertion in the active site loop (β7-α9 loop), **2**) a deletion in the β7-α9 loop, and **3**) presence of a unique Thr317 residue within the active site (Supplemental Figure 9). The active site loop insertion appears to afford additional conformational flexibility, which might impact substrate selectivity in *Pt*LYSA. Deletion in the β7-α9 loop expands the active site significantly (Supplemental Figure 9), and potentially removes steric hindrances, which could allow multiple stereoisomers to bind; the notion is consistent with the DSF analyses (Figure 6). A deletion in β7-α9 loop has also been observed in *E. coli* DAPDC (Supplemental Figure 9), which was shown to be able to bind D-lysine in the active site (PDB: 1KO0). Notwithstanding, the most remarkable adaptation in the active site of *Pt*LYSA is the replacement of the conserved arginine in the EPGR motif with a threonine (Supplemental Figure 9). Similar substitution at this residue are observed in all sequenced diatoms to date, as well as some bacterial (*Leptospira interrogans*), algal (*Ostreococcus lucimarinus*) and cryptophytes (*Guillardia theta*) lineages. The side chain of Thr317 coordinates a SO_4_, which occupies the position of PLP-PO_4_ (Figures 7 and 8). The shorter sidechain of Thr317 may relax substrate specificity and structural constraints imposed by the arginine in structurally characterized DAPDC orthologs. Importantly, each of these three active site adaptations are not specific to *P. tricornutum* and are conserved across all putative diatom LYSA proteins.

## METHODS

### Phylogenetic Analysis

Sequences for the analysis were retrieved from NCBI-nr, PhyloDB, JGI, and MarDB (https://mmp.sfb.uit.no/databases/mardb/) using DIAMOND BLAST (Buchfink, 2015). Outgroup (non-diatom) sequences were clustered on 50% identity and 80% coverage using MMseqs2 (Steinegger and Soding, 2017). Datasets were aligned by MAFFT v7.407 (Katoh and Standley 2013) using the L-INS-i refinement and a maximum of 1000 iterations and trimmed by trimAl v1.4 (Capella-Gutiérrez, 2009) allowing 70% sequences to have a gap (−gt 0.3). Maximum likelihood (ML) trees were inferred by IQ-TREE v 1.6.12 (Nguyen, 2015) using the LG+C20+F+G model and the posterior mean site frequency method (Wang, 2018), starting from a LG+F+G guide tree and employing the strategy of rapid bootstrapping followed by a “thorough” ML search with 1,000 ultra-fast bootstrap replicates.

To resolve the topology of LYSA with multiple long branches, the ML tree was compared with a Bayesian topology. The latter was inferred by PhyloBayes MPI v1.7 (Lartillot, 2013) under the CAT-GTR model most robust against long-branch attraction artifacts in three independent chains run for ~40,000 generations when they reached convergence at 0.128 maximum discrepancy. Then, burn-in of 4,000 generations (10 %) was discarded from each chain and a consensus tree was calculated from every-10^th^-tree subsamples.

### Plasmids for Gene Deletion Studies

Vectors for inhibiting the expression of *PtLYSA* via RNAi were designed using previously described plasmids (Allen et al., 2011). The plasmid backbone was linearized via PCR using primers LYS-038/039. 250-bp and 400-bp targeting fragments were amplified from the *PtLYSA* gene using primers LYS-040/041 and LYS-041/042, respectively. Loop-hairpin overlaps targeting *PtLYSA* were assembled using Gibson cloning. The resulting vectors drive loop-hairpin expression using the relatively high expression promoter *pFCPB* coupled with the terminator *tFCPA*. In a subsequent build, this vector was linearized using primers that exclude the *pFCPB* element and then assembled with *pNR* using Gibson cloning.

Plasmids for TALEN were constructed as previously described (Weyman et al., 2015) and introduced via electroporation. Plasmids for *P. tricornutum* CRISPR were built using Golden Gate-based assembly methods for CRISPR-Cas9 targeting vectors (dx.doi.org/10.17504/protocols.io.4acgsaw).

### Plasmids for Localization

*Pt*LYSA-YFP fusion constructs were built with Gateway cloning method (Siaut et al., 2007). Plasmids for episomal overexpression of fusion proteins in *P. tricornutum* were based upon *Pt*PBR1 and contained an *oriT* to allow for conjugation of the plasmids from bacteria (Karas et al., 2015). The predicted promoter and terminator for *PtLYSA* were amplified from *Phaeodactylum* gDNA and correspond to the genomic coordinate chromosome 20(+):362797-363410 for the promoter and chromosome 20(+):366255-366898 for the terminator. The nitrogen-inducible pNR promoter and native tNR terminator were amplified from *Phaeodactylum* gDNA and correspond to the genomic coordinate chromosome 20(+):362797-363410 for the promoter and chromosome 20(+):366255-366898 for the terminator. The coding sequence for the mTurquoise2 reporter protein (Goedhart et al., 2012) was amplified from plasmid L1_23 (Pollak et al., 2019) using primers LYS-09 and LYS-010 (Supplemental Information) and contained overhangs to produce a C-terminal fusion protein when translated. Plasmids were built using the Gibson assembly method and validated by Sanger sequencing.

### Plasmids for Protein Expression, DSF and Crystallography

The coding sequences of full-length, N-terminally Δ24 and Δ36 deletion constructs of *Pt*LYSA were cloned from *P. tricornutum* genomic DNA using primers described in Table 1 (Supplemental Information) and Primestar polymerase (Takara) with primers LYS13-LYS16. These fragments were sub-cloned using Gibson assembly methods into XhoI-linearized PTpBAD-CTHF vector (Brunson et al., 2018). Sequence analysis after cloning (Eurofins) revealed that some Δ24 and Δ36 clones had obtained a point mutation at the junction between the terminal F481 and thrombin cleavage linker in the vector backbone, resulting in a F481L mutation. These constructs (Δ24-*Pt*LYSA-F481L and Δ36-*Pt*LYSA-F481L) were analyzed alongside their wild-type counterparts for their level of solubility and expression.

DSF and crystallization studies utilized a DNA segment encoding *Pt*LYSA (amino acid residues 39-476) was inserted in frame into pNIC28 Bsa4 (pSGC-His) using ligation independent cloning (Gileadi et al., 2008); the gene is fused to a leader sequence encoding an TEV-cleavable N-terminal His_6_ tag. The inserted DNA sequence of the corresponding domain was sequenced (Genewiz) completely to exclude the acquisition of unwanted coding changes during DNA amplification and cloning. The resultant plasmid was transformed into *E. coli* BL21 (DE3) CodonPlus RIL (Novagen). For protein preparation, 2 L of bacterial culture (Super Broth, Teknova) supplemented with 50 μg ml^−1^ kanamycin, 100 μg ml^−1^ chloramphenicol and 100 μL L^−1^ antifoam 204 (Sigma), was grown at 37 °C in LEX 48 airlift bioreactors (Epiphyte3, Canada). After 7 hours of growth (*A*_600_ of 2), the temperature of the cultures was reduced to 20 °C for optimal protein folding, induced with addition of 0.5 mM isopropyl-β-D-thiogalactoside (IPTG; GoldBio) and incubated overnight.

### Transformation of plasmid DNA into *P. tricornutum*

The fusion constructs and control plasmids were transferred into *P. tricornutum* cells using the multi-well conjugation method (Diner et al., 2016). Briefly, plasmids were transformed into EPI-300 *E. coli* cells (Lucigen) harboring the pTA-MOB conjugation helper plasmid (Strand et al., 2014). The bacterial strains were then isolated, amplified and mixed with diatom cells to allow for conjugation of plasmids according to published methods (Diner et al., 2016). Diatom exconjugants were selected on _1/2_L1 1% agar medium supplemented with 20 μg mL^−1^ phleomycin (Gold Biolabs) at a diel cycle of 14:10 at ~50 μE ms^−1^ light intensity.

### Microscopy

All confocal images were captured using a Leica TCS-SP5 confocal microscope (Leica Microsystems). mT2 was detected by excitation with a 458 nm laser and an emission range of 465-510 nm. Chlorophyll autofluorescence was also excited with a 458 nm laser and detected with an emission range of 680-712 nm.

### Small-scale expression of *Pt*LYSA

Clones from BL21 *E. coli* (New England Biolabs) transformation plates were picked and cultured overnight in 5 mL LB broth supplemented with 10 μg mL^−1^ tetracycline at 30 °C and 220 rpm shaking. On the day of testing, these cultures were used to seed 10 mL cultures of Terrific Broth supplemented with 10 μg mL^−1^ tetracycline at a 1:20 dilution. The cultures were then incubated for 4 hours at 30 °C, at which time they were then moved to 18 °C for an additional 60 minutes to cool. L-arabinose was added to the cultures to a final concentration of 0.5% and the cells allowed to incubate overnight at 18 °C. Cultures were pelleted the following day and evaluation of protein expression was carried out as described (dx.doi.org/10.17504/protocols.io.bkhgkt3w). Eluates and insoluble fractions were analyzed via polyacrylamide electrophoresis and gels stained with Coomassie Brilliant Blue G-250 solution for visualization of proteins.

### Purification of *Pt*LYSA for biochemical analysis

For large-scale purification, 1 L of the Δ36-*Pt*LYSA-F481L BL21 cell line was cultured and induced as described above. Cell pellets were sonicated in 500 mM NaCl, 20 mM Tris-HCl pH 8.0, 10% glycerol and purified using FPLC and a 5 mL HisTrap FF column. Large-scale protein purification from *E. coli* was performed on an ÄKTApurifier instrument (GE Healthcare) with the modules Box-900, UPC-900, R-900 and Frac-900 with all solvents filtered through a nylon membrane 0.2 μm GDWP (Merck) prior to use. Final eluates were pooled and resuspended in 20 mM HEPES pH 8.0, 300 mM KCl, and 10% glycerol. Concentration was determined by method of Bradford.

### Decarboxylation Assay

Enzyme assays for HPLC analysis were conducted in 50 mM HEPES pH 8, 200 mM KCl, and 10% glycerol, at 100 μL scale. Substrates (DAP, ornithine and arginine) were added at 1 mM concentration and cofactor PLP was added at 50 μM concentration. Assays were started with the addition of ~50 μM *Pt*LYSA protein and allowed to incubate overnight (12-18 hours), after which reactions were quenched upon addition of assay components for Marfey’s derivatization.

### Marfey’s derivatization and RP-HPLC analysis

Marfey’s derivatization was carried out by the addition of 20 μL concentrated Na_2_CO_3_ (approx. 10% w/v) and 100 μL of freshly prepared 1% w/v 1-fluoro-2,4-dinitrophenyl-5-L-alanine amide (L-FDAA) in acetone to 50 μL of the *Pt*LYSA reactions described above. For the generation of standards and H_2_O blanks, 50 μL of 1 mM aqueous amino acid stock solutions or Milli-Q H_2_O was added in place of the *Pt*LYSA reaction mix. Marfey’s derivatization reactions were allowed to incubate for 90 minutes at 37 °C before quenching with 25 μL 1N HCl and adding 5 μL Milli-Q H_2_O to bring the total volume to 200 μL. Reactions were centrifuged (18,000 × *g*, 10 min) and 10 μL of the clarified supernatant was injected for analytical reverse-phase HPLC (Agilent Technologies 1200 series, Phenomenex Luna 5u C18(2), 4.6 × 150 mm) at a flow rate of 1 mL min^−1^ using the following method: 5% B (5 min), 5 – 95% B (20 min), 95 – 5% B (1 min), 5% B (4 min), where A = 0.1% aqueous trifluoroacetic acid, and B = 0.1% trifluoroacetic acid in acetonitrile (Brunson et al., 2018). Reaction products were monitored by UV-detection at 340 nm.

### Protein purification of *Pt*LYSA for DSF and Crystallization

Cells were harvested by centrifugation at 6,500 × g and suspended in buffer containing 20 mM HEPES (pH 7.5), 500 mM NaCl, 20 mM imidazole, 0.1% IGEPAL, 20% sucrose, 1 mM β-mercaptoethanol (βME). Cells were disrupted by sonication and cell-debris was removed by centrifugation at 45,000 × g. The supernatants were applied to a chromatography column packed with 5 ml His60 superflow resin (Clontech) that had been equilibrated with buffer A (20 mM HEPES pH 7.5, 20 mM imidazole, 500 mM NaCl, 1 mM βME). The column was washed with buffer A and the His_6_ tagged *Pt*LYSA was eluted with buffer B (20 mM HEPES pH 7.5, 350 mM NaCl, 250 mM imidazole, 1 mM βME). The N-terminal His_6_ tags from the protein was removed by overnight digestion at 4 °C with the TEV protease at a 1000:1 (w/w) ratio of *Pt*LYSA:TEV. The tag-free protein was then separated from the His_6_ tag and TEV protease by a Superdex 200 (16/60) size exclusion chromatography column equilibrated with buffer containing 20 mM HEPES pH 7.5, 150 mM NaCl, and 5 mM DTT. Potential peak fractions containing *Pt*LYSA were assessed by SDS-PAGE, pooled concentrated to 30 mg ml^−1^ using a 30 kDa Amicon Ultra-15 centrifugal filter device (Millipore). For crystallization experiments, the tag-free protein was purified using a Superdex 200 (16/60) gel filtration column in a buffer containing 20 mM Na-Acetate pH 5.5, 350 mM NaCl, and 1 mM βME. Fractions containing the peak of the protein were pooled and concentrated to 20 mg/ml and buffer exchanged to Na-Acetate (pH 5.5), 150 mM NaCl, and 10 mM DTT using a 30 kDa Amicon Ultra-15 centrifugal filter device (Millipore). Purified *Pt*LYSA appeared to be bright yellow, suggesting the presence of co-purified cofactor, PLP from *E. coli* host.

### Differential Scanning Fluorimetry

Potential ligands (10 mM) stocks were prepared in H_2_O and stored at −20 °C, and diluted to 1 mM working stock in buffer containing 20 mM HEPES, pH 7.5, 150 mM NaCl and 5 mM DTT (DSF buffer). Final DSF reaction mixtures (20 μl final volume) contained 100 μM *Pt*LYSA, 500 mM each ligand, and 5× SYPRO Orange (Invitrogen) in DSF buffer and were distributed in a 384-well PCR plate (Applied Biosystems) with 5 control wells (purified *Pt*LYSA) and 5 wells with *Pt*LYSA-ligand complexes indicated in Figure 6. The fluorescence intensities were measured using an Applied Biosystems 7900HT fast real-time PCR system with excitation at 490 nm and emission at 530 nm. The samples were heated from 25 to 99 °C at a rate of 3 °C min^−1^. The midpoint of the unfolding transition (T_m_) was obtained from fitting the melting curve to a Boltzmann equation (Niesen et al., 2007). ΔT_m_ for each specific ligand was calculated as the difference of the T_m_ values measured with or without a ligand (average of 5 measurements).

### Protein crystallization and crystal harvesting

Initial crystallization screening of *Pt*LYSA was performed using 800 nl (protein:mother liquor=1:1) sitting drops at a concentration of 12 mg/ml (255 μM) co-crystallized with D-lysine (1 mM) with a Crystal Gryphon (Art Robbins Instruments) and utilizing MCSG sparse matrix crystallization suite (Microlytic). Crystals were eventually obtained in sitting drops by vapor diffusion against well solutions containing 2.0 M Ammonium sulfate and 0.1 M Bis-Tris pH 6.5. Crystals appeared after 12 days at 19 °C and reached maximum dimensions of 15 × 30 × 30 μm^3^ in another week. For data collection, crystals were cryo-preserved by addition of 30% glycerol to the mother liquor prior to flash-cooling in liquid nitrogen.

### Data collection and processing, structure determination, model building, refinement and analysis

Data were collected with an Eiger 9M detector, with a wavelength of 0.98 Å, on the ID-17-1 (AMX) beamline at the National Synchrotron Light Source-II, Brookhaven National Laboratory (Table 2). Data from a single crystal were integrated and scaled using AIMLESS (Winn et al., 2011). Diffraction was consistent with the orthorhombic space group P2_1_2_1_2_1_ (a=85.8, b= 86.3, c=127.2 Å) and extended to a resolution of 2.78 Å, with two molecules in the asymmetric unit (protomer 1/chain A and 2/chain B). Initial phases were determined by molecular replacement with PHASER (McCoy et al., 2007) using refined coordinates of the *E. coli* DAPDC (PDB: 1KNW, Seq ID 27%) as the search model. After determining the phases an atomic model was built into the density using the automated model building program BUCCANEER (Cowtan, 2006) and manually inspected using COOT (Emsley and Cowtan, 2004). The model was refined with REFMAC5 (CCP4, 1994) and PHENIX (Liebschner et al., 2019). For structural interpretation of *Pt*LYSA protomer 1 has been used. Analyses of the structures were performed in COOT and MOLPROBITY (Chen et al., 2010). Crystallographic statistics and RCSB accession codes are provided in Table 2.

## Accession Numbers

This study did not generate new genomic data. Accession numbers for genes described in this work include *P. tricornutum LYSA*, J21592; *DAPA*, J11151; *DAPB*, J4025; *DAPL*, J22909; *DAPF*, J34852.

## Supplemental Data

**Supplemental Figure 1**. Phylogenetic analysis of the putative diatom lysine biosynthesis pathway components.

**Supplemental Figure 2**. Diel expression of lysine pathway genes from Table 1 from transcriptomic analysis.

**Supplemental Figure 3**. Reduction of *Pt*LYSA protein content in *P. tricornutum* cells via RNAi results in a lag in growth rate.

**Supplemental Figure 4**. Sanger sequencing of CRISPR *P. tricornutum* exconjugants.

**Supplemental Figure 5**. Small-scale solubility testing of *Pt*LYSA-CTHF constructs in *E. coli* BL21 cells.

**Supplemental Figure 6**. *Pt*LYSA does not act upon L-ornithine.

**Supplemental Figure 7**. Structure based primary sequence analyses.

**Supplemental Figure 8**. Dimer interface of *Pt*LYSA.

**Supplemental Figure 9**. The LEEAA segment of *Pt*LYSA maintains integrity of the active site.

## ACKNOWLEDGEMENTS

This research is supported by the *Department of Energy*, Office of Biological and Environmental Research (*BER*), grant DE-SC0018344.

## AUTHOR CONTRIBUTIONS

V.A.B., J.K.B., A.G., M.A.M, S.M.K.K., and C.L.D. designed research. V.A.B., J.K.B., A.G., M.A.M., E.A.G., Z.F., J.B., S.M.K.K., and C.L.D. performed research. V.A.B., J.K.B., A.G., M.A.M., E.A.G., and C.L.D. contributed new reagents/analytic tools. V.A.B., J.K.B., A.G., M.A.M., Z.F., S.M.K.K., B.S.M., A.E.A., S.C.A., and C.L.D. analyzed data. V.A.B., J.K.B., A.G., M.A.M., S.C.A., and C.L.D. wrote the article.

---

**Figure S1.**
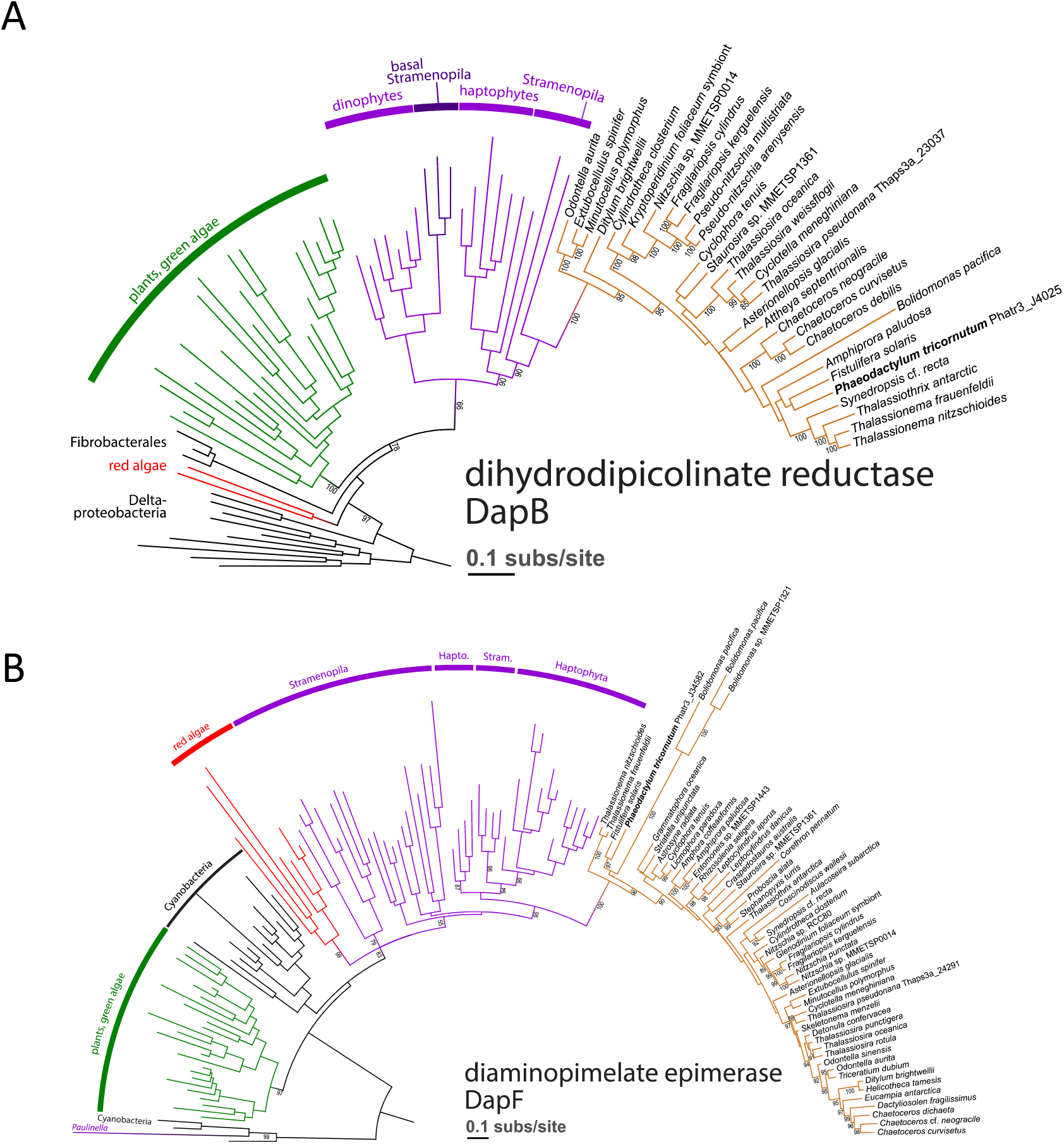
Phylogenetic analysis of the putative diatom lysine biosynthesis pathway components. **(A)** ML phylogeny of dihydrodipicolinate reductase *DapB*. With moderate support, plants and green algae are more related to complex algae, including diatoms. The *DapB* homolog in this clade is likely related to delta-proteobacteria or Fibrobacterales (note the long branch of the latter group). In this schematic, bootstrap support is shown for critical branching points of the outgroups, while >85 branch support is shown in the Bolidophyceae-diatom clade. **(B)** ML phylogeny of diaminopimelate epimerase *DapF*. This enzyme is apparently derived from cyanobacteria and in plants and green algae, red algae, stramenopiles and haptophytes constitutes a monophyletic clade, with red algae being moderately supported as the donor of the complex algal homolog. *Paulinella* are rhizarian algae that established a primary endosymbiotic relationship with a cyanobacterium independently on other algal lineages, as apparent from the figure. In this schematic, bootstrap support is shown for critical branching points of the outgroups, while >85 branch support is shown in the Bolidophyceae-diatom clade.

**Figure S2.**
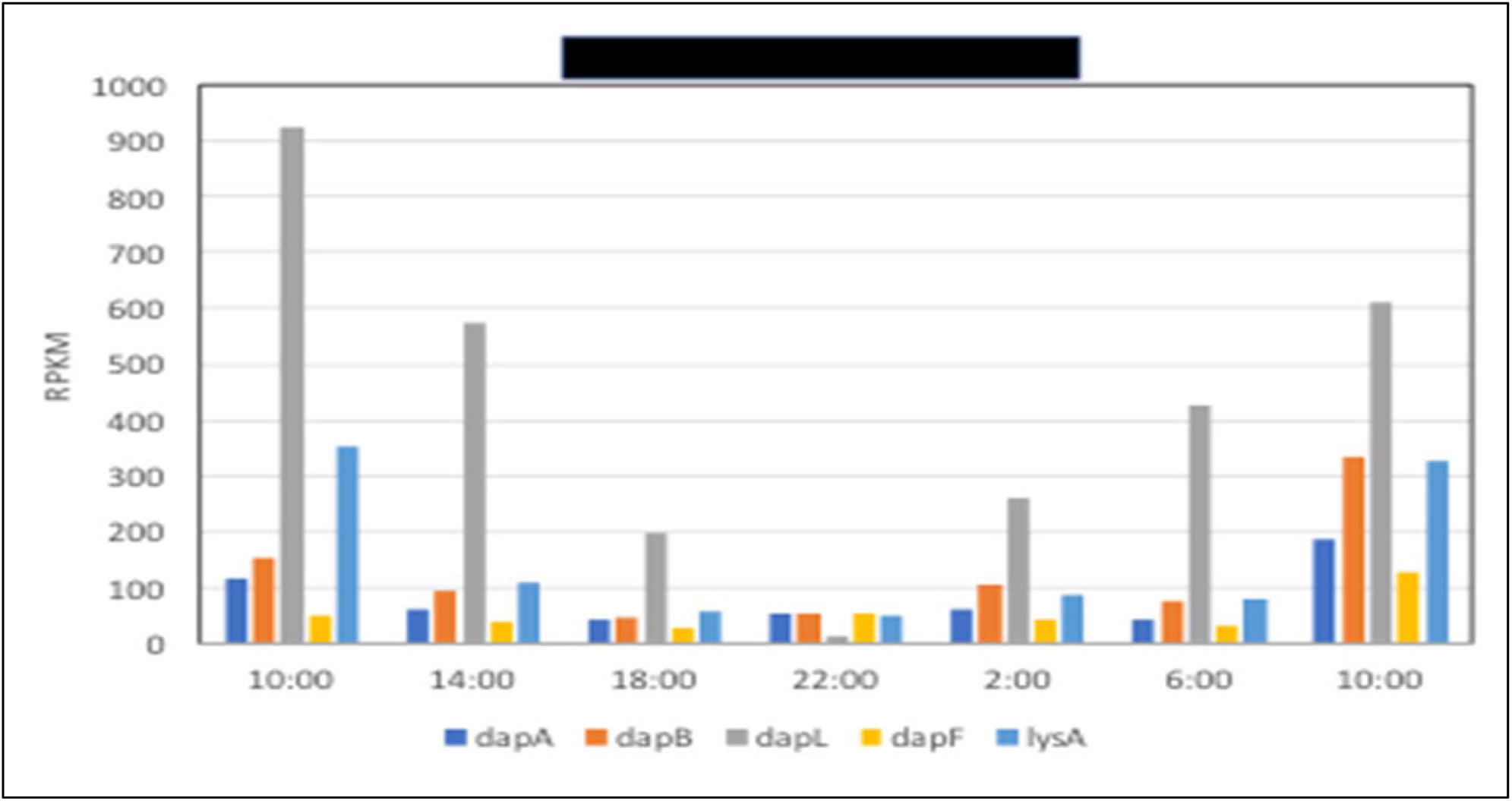
Diel expression of *P. tricornutum* lysine pathway genes from Table 1 from transcriptomic analysis. The black bar above represents the dark portion of the day. Adapted from Smith et al., 2016.

**Figure S3.**
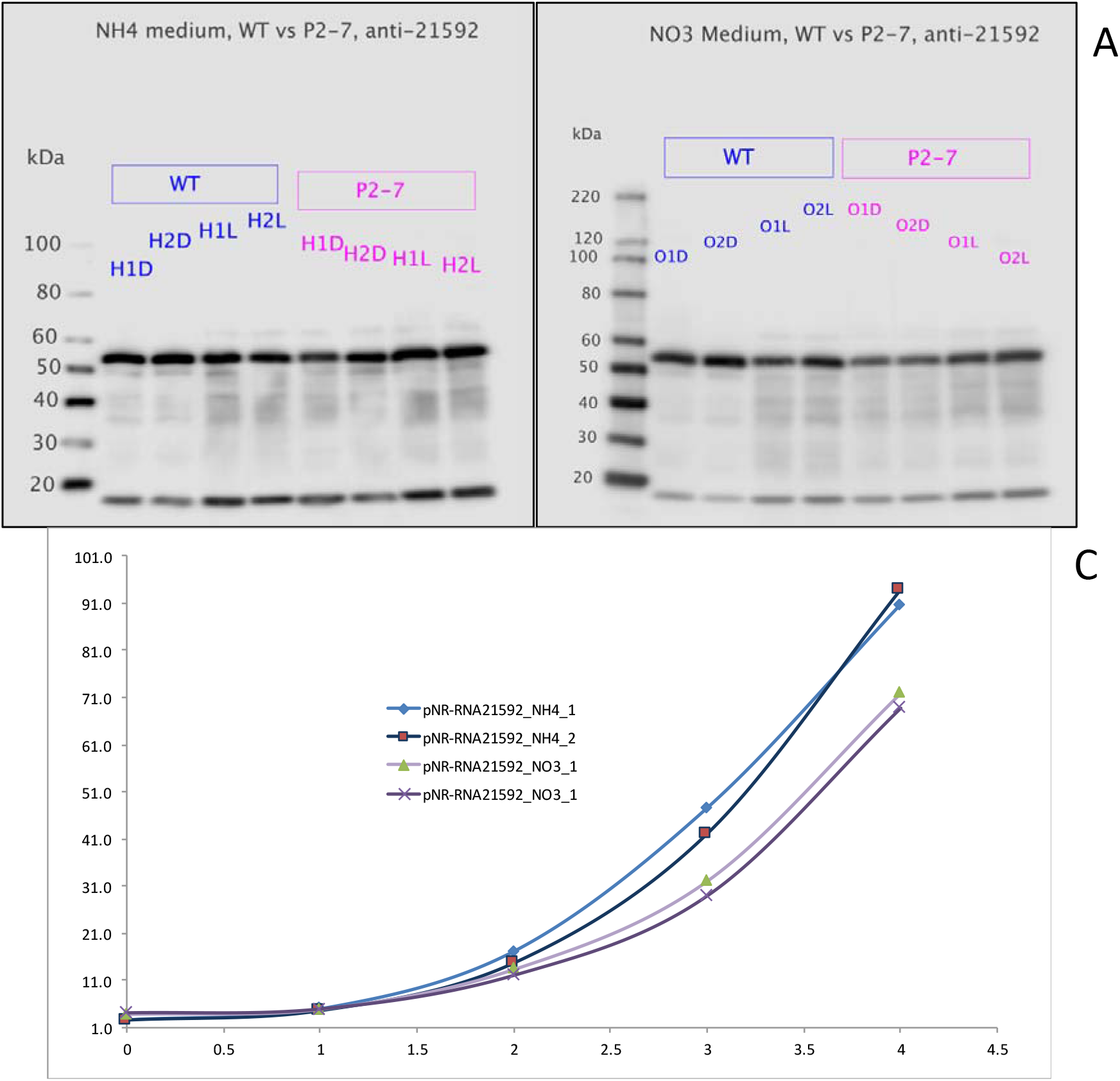
Reduction of *Pt*LYSA protein content in *P. tricornutum* cells via RNAi results in a lag in growth rate. **(A)** Western blot of PtLYSA protein in uninduced RNAI conditions (NH4-repressed) comparing wild-type (WT) versus transgenic cell line (P2-7) **(B)** Western blot of PtLYSA protein in induced RNAI conditions (NO3-active) comparing wild-type (WT) versus transgenic cell line (P2-7) **(C)** RNAI-induced vs-uninduced growth curve of P2-7 cell line in duplicate. X-axis, time in days; Y-axis, chlorophyll content of batch cultures measured via fluorimetry

**Figure S4.**
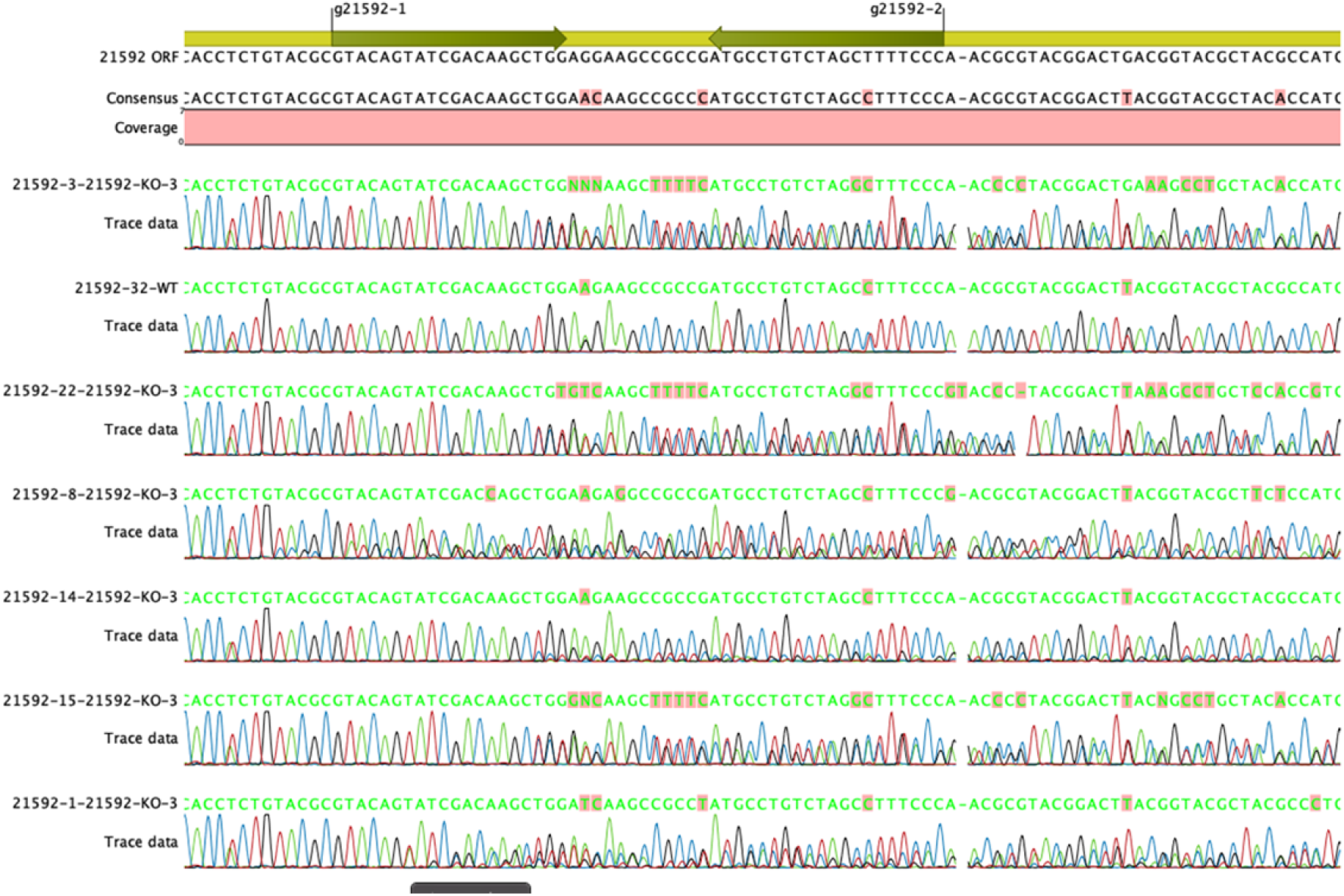
Sanger sequencing of CRISPR *P. tricornutum* exconjugants. Green arrows in schematic at top represent the targeted sites within *PtLYSA* gene and red bar represents expected 18-bp deletion upon editing with two sgRNAs.

**Figure S5.**
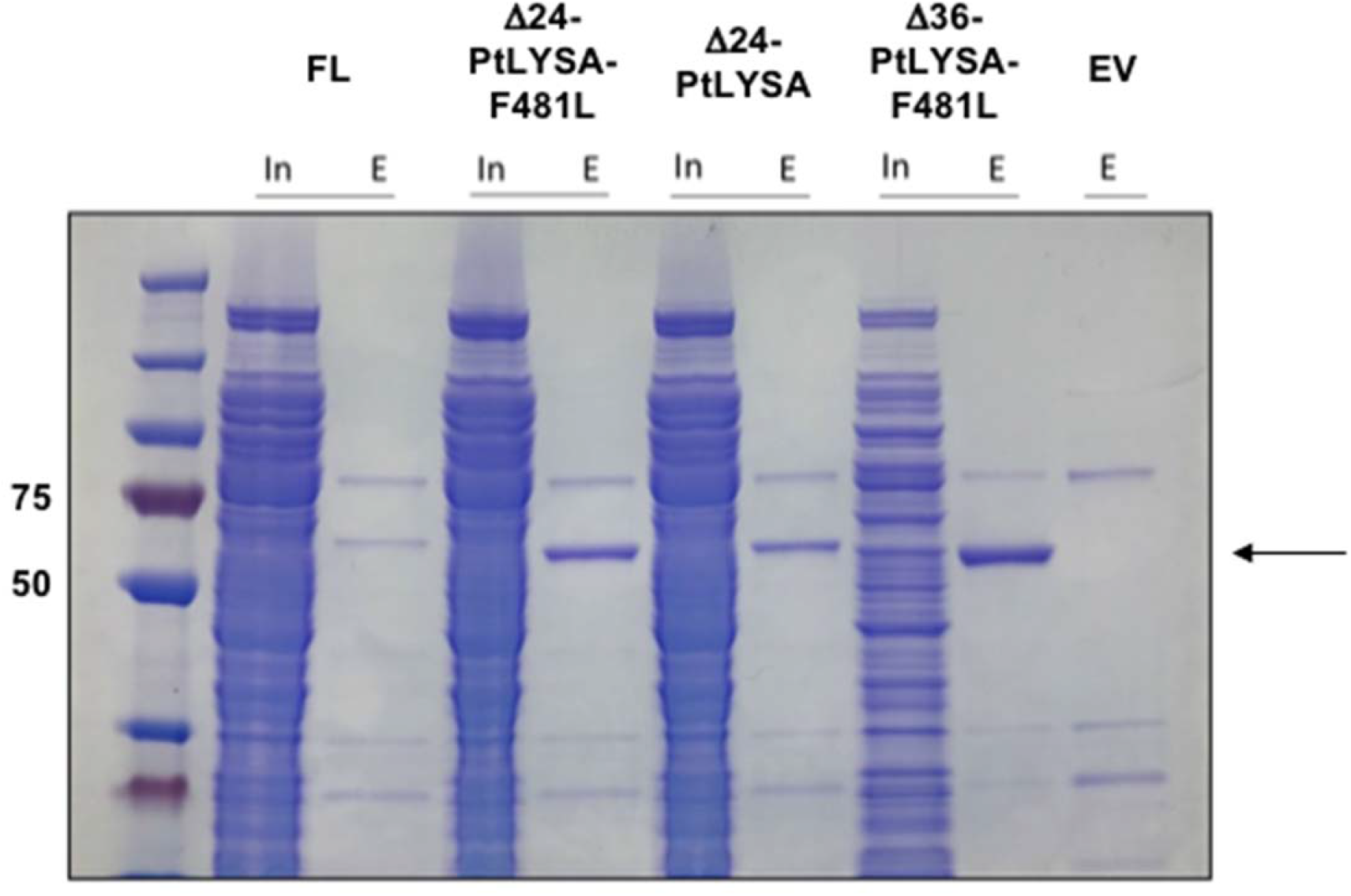
Small-scale solubility testing of *Pt*LYSA-CTHF constructs in *E. coli* BL21 cells. Protein expression was induced with 0.5% L-arabinose in Terrific Broth at 30 °C for 16 hours. The arrow indicates the Δ36-*Pt*LYSA-F481L-CTHF construct employed for biochemical analyses. In, clarified input; E, eluate; FL, full-length protein; EV, empty vector control.

**Figure S6.**
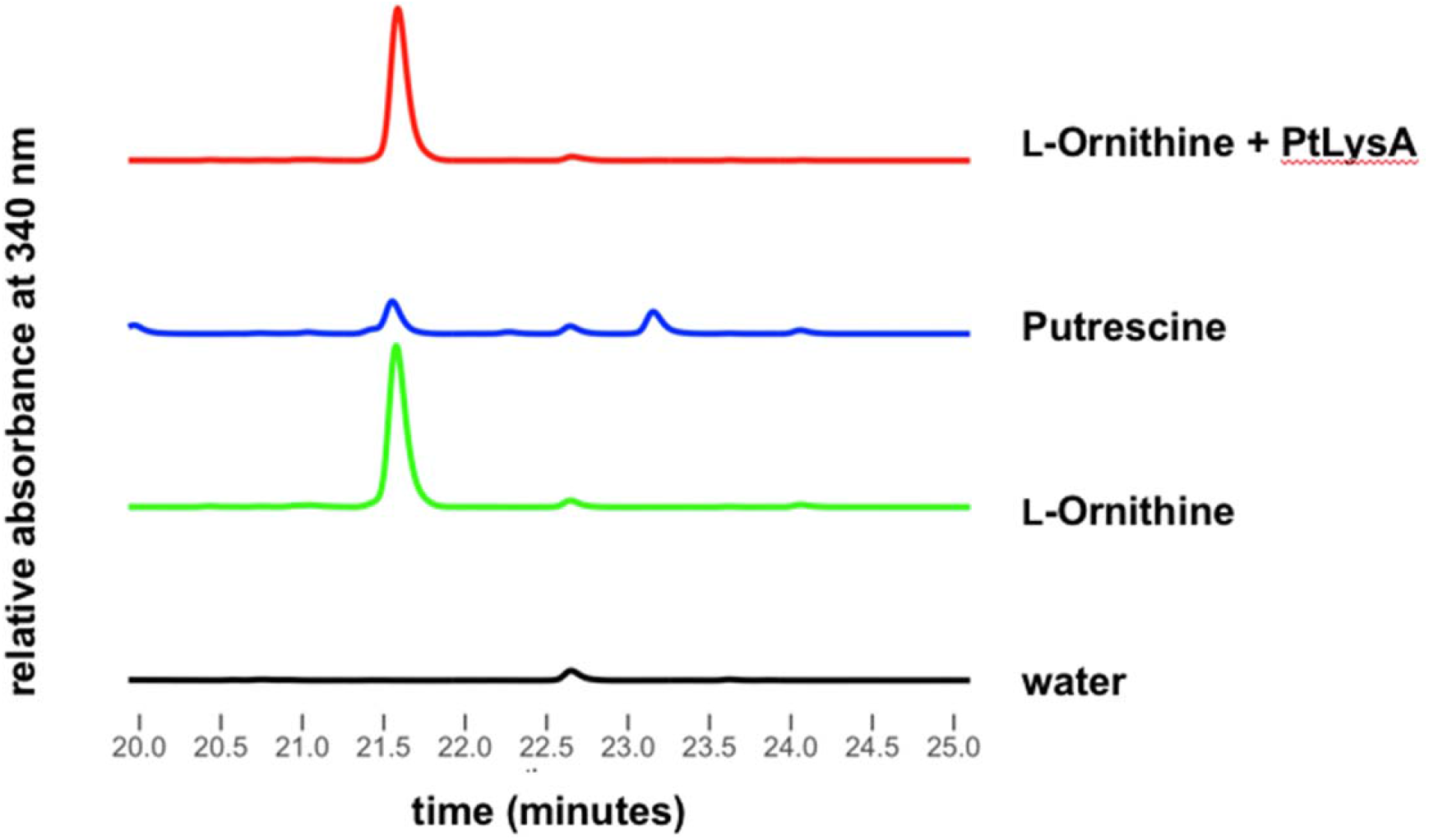
*Pt*LYSA does not act upon L-ornithine. RP-HPLC (λ = 340 nm) analyses of L-FDAA (Marfey’s) derivatized *Pt*LYSA reactions with L-ornithine and comparison to similarly derivatized ornithine (time ~21.5 min) and putrescine (time ~23.2 min) standards. Substrates for enzyme reactions were added at 1 mM concentration and similarly 1 mM of each standard was used for subsequent derivatization.

**Figure S7.**
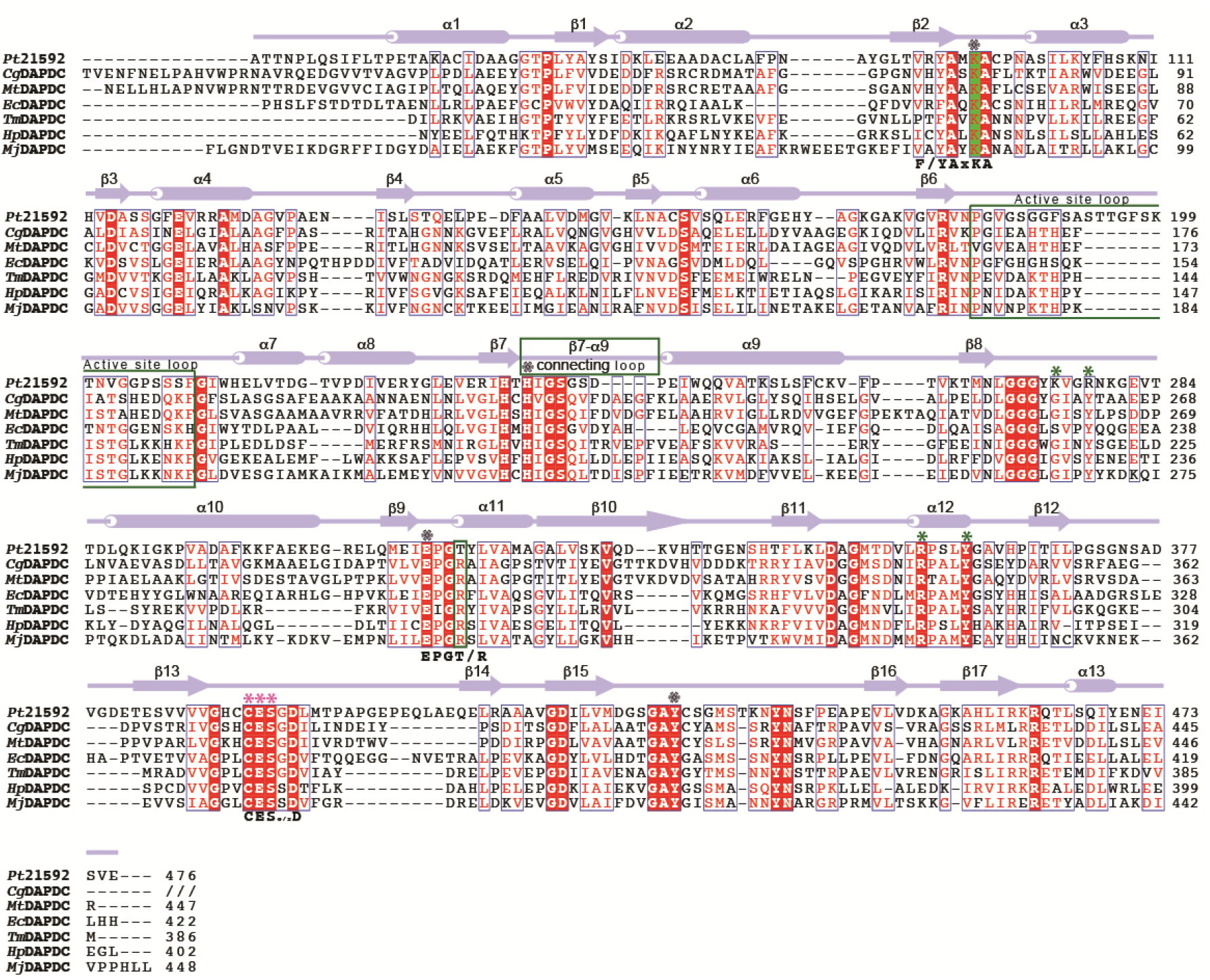
Structure based primary sequence analyses. Aligned amino acid sequences for *Pt*LYSA and DAPDC from *C. glutamicum* (PDB: 5X7M), *M. tuberculosis* (PDB: 2O0T), *E. coli* (PDB: 1KNW), *T. maritima* (PDB: 2YXX), *H. pyroli* (PDB: 2QGH) and *M. jannaschii* (PDB: 1TUF). *Pt*LYSA secondary structure elements (helices as cylinders; strands as arrows) are color coded as in Figure 6 above the sequence. Alignment gaps indicated by (−). Sidechain identity/similarity is denoted by red shading (conserved in all) or red letter (conserved in most). Location of Thr317, the β7-α9 connective and active site loops are outlined. Conserved lysine that forms Schiff’s base with PLP highlighted in green shade. Light-blue and light-pink asterisks above the alignment indicate conserved active site residues supplied in *cis* and *trans*, respectively. Green asterisks above denote *Pt*LYSA residues adjacent to putative D-lysine. Three key binding motifs are indicated below the alignment.

**Figure S8.**
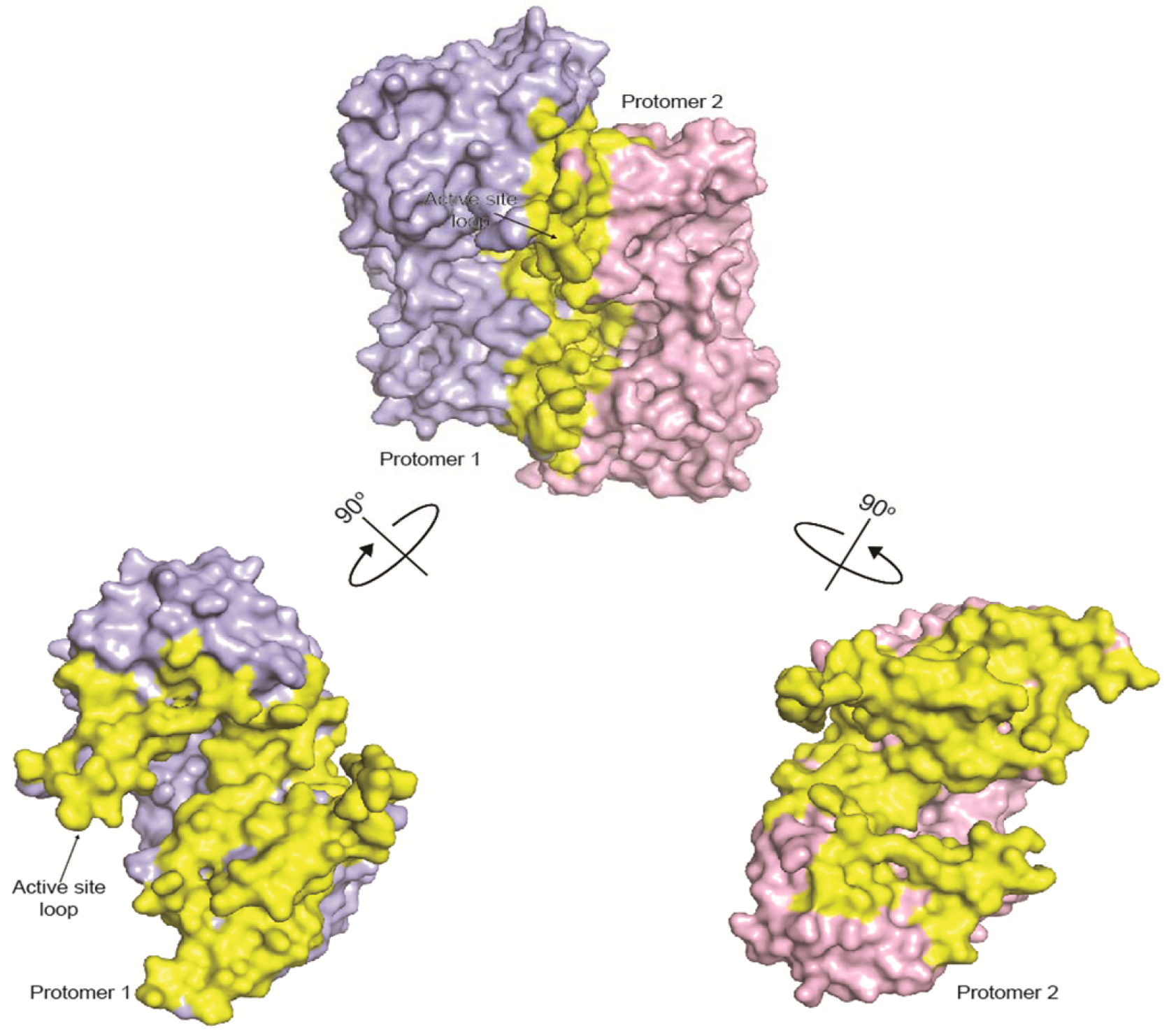
Dimer interface of *Pt*LYSA. Structure of *Pt*LYSA is shown in surface representation and the protomers are color coded as in Figure 6. The interface was mapped using 1.4 Å probe radius and colored in yellow. Position of the active site loop in protomer 1 is indicated by arrows and labeled.

**Figure S9.**
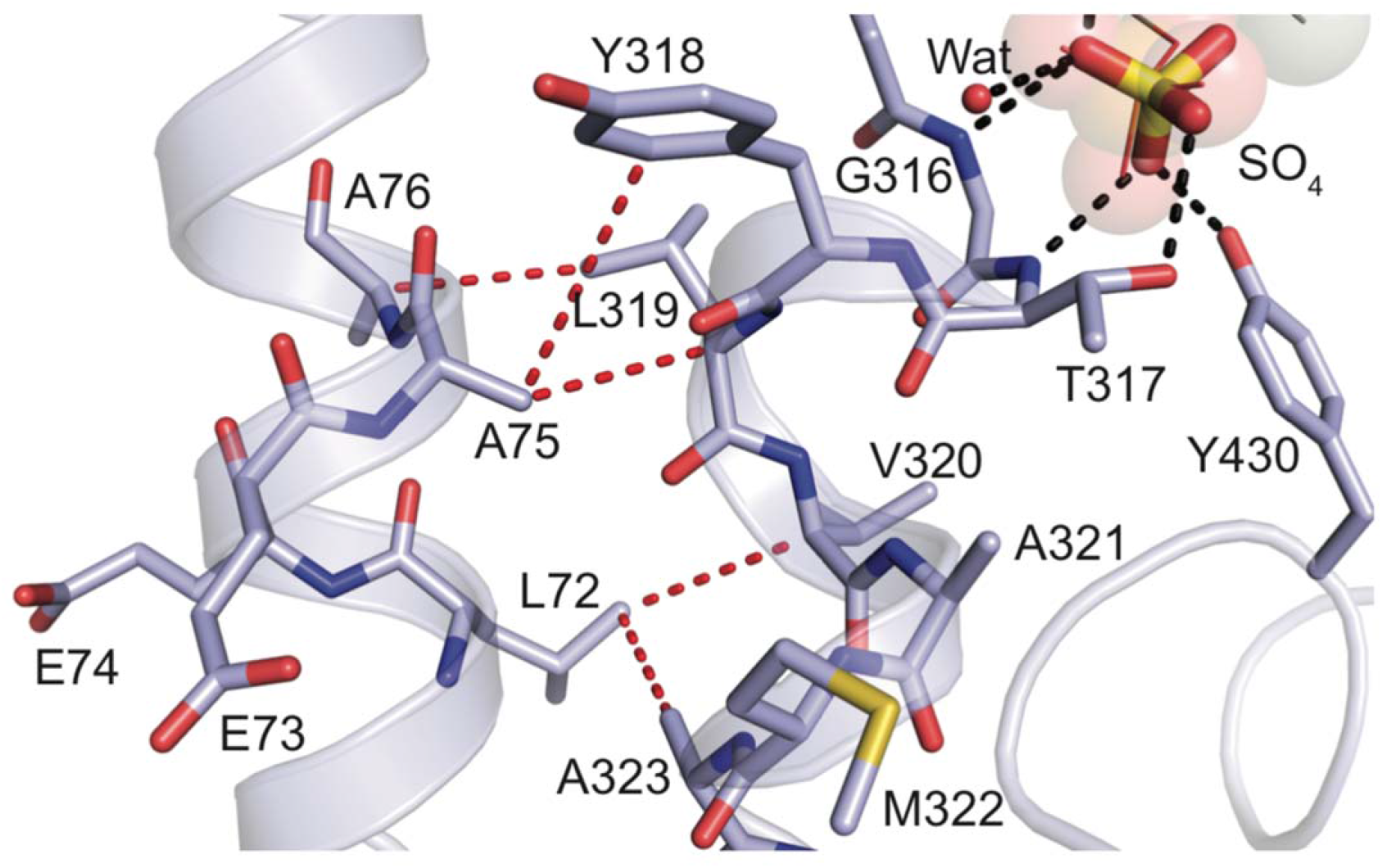
The LEEAA segment of *Pt*LYSA maintains integrity of the active site. A view of the interactions between the LEEAA (part of helix a6; aa 72-76) and the segment that demarcates the active site in protomer 1 (chain A; color coded as in Figure 6). Stick models of the amino acids that comprise the interface and the active site are shown; the bound SO_4_ and water in the active site are shown in stick representation and red sphere, respectively. Part of PLP (transparent spheres and lines) was modeled into the active site based on its position in the *C. glutamicum* structure (PDB: 5X7M) to denote the potential location and orientation of the co-factor in *Pt*LYSA. Potential hydrogen bonds and van der Waals interactions are represented as black and red dashed lines, respectively. The interaction of the LEEAA segment at the interface highlights its potential role in maintaining active site integrity.

